# Spatiotemporal solidification of α-synuclein inside the liquid droplets

**DOI:** 10.1101/2021.10.20.465113

**Authors:** Soumik Ray, Debdeep Chatterjee, Semanti Mukherjee, Komal Patel, Jaladhar Mahato, Sangram Kadam, Rakesh Krishnan, Ajay Singh Sawner, Manisha Poudyal, G Krishnamoorthy, Arindam Chowdhury, Ranjith Padinhateeri, Samir K. Maji

**Affiliations:** Department of Biosciences and Bioengineering, IIT Bombay, Powai, Mumbai-400076, India; Department of Chemistry, IIT Bombay, Powai, Mumbai 400076, India; Department of Biotechnology, Anna University, Chennai 600025, India

**Keywords:** Liquid-liquid phase separation, liquid-to-solid transition, α-synuclein.

## Abstract

Liquid-liquid phase separation (LLPS) and subsequent liquid-to-solid transition is implicated in membraneless organelles formation as well as disease associated protein aggregation. However, how liquid-to-solid transition is initiated inside a liquid droplet remains unclear. Here, using studies at single droplet resolution, we show that liquid-to-solid transition of α-synuclein (α-Syn) liquid droplets is associated with significant changes in the local microenvironment as well as secondary structure of the protein, which is prominently observed at the center of the liquid droplets. With the ageing of liquid droplets, the “structured” core at the center gradually expands and propagates over entire droplets. Further, during droplet fusion, smaller, homogeneous droplets progressively dissolve and supply proteins to the larger, heterogeneous droplets containing solid-like core at their center. The present study will significantly help to understand the physical mechanism of LLPS and liquid-to-solid transition in biological compartmentalization as well as in protein aggregation associated with human neurodegenerative disorders.

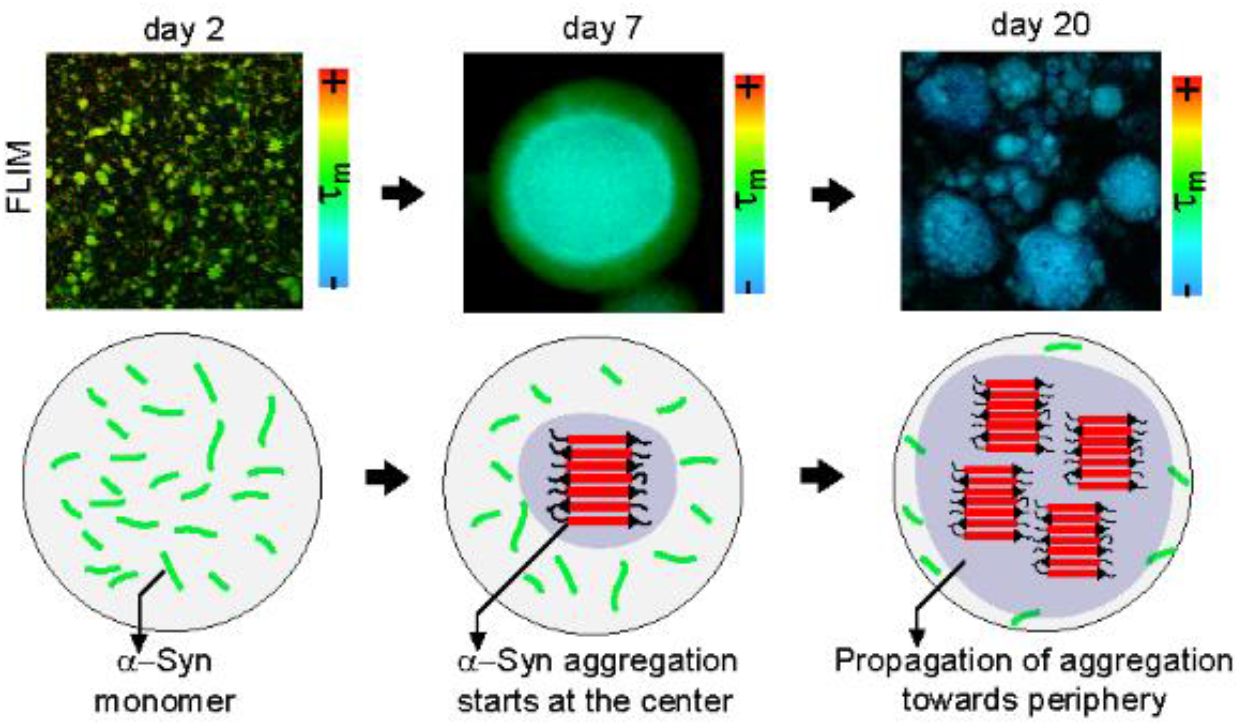

## Introduction

Liquid-liquid phase separation (LLPS) by various biomolecules such as protein and nucleic acids has been shown to be crucial for the formation of membrane-less organelles to compartmentalize biomolecules in liquid-like condensates^1–11^. The LLPS is involved in multiple cellular functions such as concentrating cellular components into organizational hubs^1–10, 12–17^, enriching component molecules for specific biochemical reactions with regulated kinetics and high specificity^18–25^ and sequester proteins in response to specific physico-chemical stimuli^26–30^. The major advantage of membrane-less organelle formation is their ability to assemble, disassemble, fuse and exchange their constituents with the surrounding according to the functional requirement of the cell^1, 4, 31–33^. Proteins with intrinsically disorder regions (IDRs) as well as closely related low complexity domains (LCDs) have been shown to undergo transient, multivalent interactions responsible for biomolecular condensation^12, 34–40^. Apart from various important functional roles of LLPS, liquid droplets formed by many IDR containing proteins have shown to undergo liquid-to-solid transition during time^12, 13, 41–46^. Liquid-to-solid phase transition is a consequence of strong interaction between protein molecules owing to their entanglement, rapid vitrification and amyloid fibril formation (and network formation through lateral association)^22^. Once solidified, the droplets can no longer fuse, deform or exchange their constituents with the surroundings^12, 41–46^.

Notably, liquid-to-solid transition of LLPS is often associated with many biological functions of the host organism^7, 47–49^. Interestingly, such liquid droplets and bimolecular condensates could contain a solid fraction as a sub-structure^22^. However, the ratio of solid and liquid fractions could be actively regulated by cells to optimize functionality of such heterogeneous liquid droplets^22^. Apart from biological functions, liquid-to-solid transition is also associated with toxic oligomer and fibril formation associated with human diseases^50–53^. For example; LLPS of FUS^41, 42^ and hnRNPA1^12^ is important for formation and assembly of stress granules; but deregulation of LLPS of these proteins leads to aberrant solidification and fibril formation in amyotrophic lateral sclerosis (ALS). The balance between functional versus aberrant LLPS of proteins could also be modulated by factors such as *in cell* environment, disease associated mutations and post-translational modifications in proteins^12, 36, 39, 41, 42, 54–57^. Natively unstructured proteins associated with neurodegenerative disorders have also been reported to undergo LLPS and liquid-to-solid transition^41–44^. In this direction, recently, we showed that α-synuclein (α-Syn) could also phase separate into liquid droplets both *in vitro* and *in cell*^55^. Interestingly, these droplets act as nucleation sites for α-Syn aggregation and undergo a viscoelastic transition with time (from liquid-like to solid-like). The droplets eventually mature into an amyloid fibril containing hydrogel *in vitro*^55, 58, 59^.

Although liquid-to-solid transition is prevalent in biological LLPS, but the mechanism of liquid-to-solid transition and the location where solidification begins is unknown. Rigidification within the droplets could start at the center, periphery, or could be completely at random locations. In this study, we used fluorescence lifetime imaging (FLIM), single point Fourier transform infra-red microscopy (FTIRM) imaging, spatially resolved Förster resonance energy transfer (FRET) microscopy and fluorescence recovery after photobleaching (FRAP) to unveil how liquid-to-solid transition is initiated inside α-Syn liquid droplets. Our data revealed that a solid-like core forms at the center of liquid droplets, which gradually progresses towards the periphery until the entire droplet is rigidified. We further show that the formation of the solid-like core is due to secondary structural transition of α-Syn into β-sheet rich amyloid fibrils via α-helix rich intermediate formation. Furthermore, analyses of droplet growth via fusion revealed that the droplet containing a structured core grows in the expense of smaller droplets without structured core. Together, our findings prove that liquid-to-solid transition of α-Syn LLPS follows a specific pattern (center◊periphery), which could be a generic phenomenon for many natively unstructured proteins containing LCDs/IDRs that undergo such aberrant phase transitions.

## Results

### Fluorescence life-time imaging (FLIM) shows changes in α-Syn microenvironment at the center of liquid droplets, which extends towards the periphery during liquid-to-solid transition

To delineate the changes in protein property during liquid to solid transition, we phase separated α-Syn using a condition^55^ (200 μM α-Syn + 10% (w/v) PEG-8000 in 20 mM sodium phosphate buffer, pH 7.4), where α-Syn phase separation was slow compared to high salt conditions^60^. Immediately after their formation and during liquid-to-solid transition, we employed FLIM studies to delineate spatiotemporal change of protein microenvironment inside the droplets. Important to note, that FLIM studies can be used to delineate the mean fluorescence lifetime (τ_m_) of proteins independent of their concentrations (or fluorophore concentration)^61, 62^. Moreover, due to the involvement of central hydrophobic domain (NAC domain) of α-Syn (61-95 amino acids) in LLPS^55^ as well as in aggregation^63^; we used 74-Cysteine-α-Syn mutant (74C-α-Syn) labeled with fluorescein-5-maleimide^55^ for FLIM measurements (**Supplementary Fig 1-2)**. Fluorescein labeled α-Syn showed observable liquid droplets at day 2 (d2)^55^ with frequent fusion event, which was visible under fluorescence microscope, suggesting their liquid-like nature (**Fig 1a** and **Supplementary Fig. 2**). For FLIM imaging, the central plane along the Z-axis (depth) was chosen (**Fig 1b, *Left panel***). To probe the spatial dependence, we collected lifetime data at the center (C) and on three concentric circles (inner circle, IC; outer circle, OC and periphery, P) equally spaced from each other (**Fig 1b, *Right panel***). In addition, we acquired similar data from equally spaced focal planes (plane 1→3) along the Z-axis to analyze the droplets along their depth (Z-axis) (**Fig 1c).** From the FLIM images (**Fig 1d, *Left panel***), time-resolved fluorescence intensity decay profiles from all four regions (C, IC, OC and P) on the central plane were acquired (**Fig 1d, *Middle panel***), which could be satisfactorily fit to biexponential decay functions (**Supplementary methods**) to obtain two lifetimes—one long fluorescence lifetime(Δ_m_) and one short fluorescence lifetime (λ_m_). Interestingly, our observations show that at d2, there were no significant differences between the two lifetimes (Δ_m_, λ_m_) across different regions (C, IC, OC and P) of the droplets (**Fig 1d, *Right panel***). Therefore, the local microenvironment of α-Syn was identical throughout the droplets at early stages of LLPS (d2). In contrast, similar analysis confirmed a significant difference in the Δ_m_ for different regions at d5 (C bearing the lowest Δ_m_=2.74 ns and P bearing the highest Δ_m_=7.06 ns). IC and OC showed Δ_m_ of 3.03 and 5.56 ns, respectively (**Fig 1d, *Right panel***). Similar trend was observed for d10 droplets. Strikingly, d15 droplets showed an overall decrease in the Δ_m_ values across all positions on the droplet (Δ_m_ of C, IC, OC and P were calculated to be 1.7 ns, 2.03 ns, 2.52 ns and 2.97 ns, respectively). Δ_m_ was further decreased for d20 droplets. However, the data unveiled no significant differences with respect to different regions of the droplets at d20 (**Fig 1d, *Right panel***). Additionally, the λ_m_ was found to be similar (∼1.00 ns) for different regions and no change was noted during the timescale (d5-d20) of our experiment. The amplitudes of λ_m_ and Δ_m_ were almost equal (50%) in most of the cases. Since we did not know the origin of the two lifetimes, we calculated the mean fluorescence lifetimes (τ_m_) as a comparative measure of our analyses **(Fig 1e)**. Our results revealed that there is no significant difference between τ_m_ (∼4 ns; denoted with green pixels) across different regions of the droplets at d2 **(Fig 1e-f),** which is similar to soluble, monomeric protein (τ_m_=4.1 ± 1 ns) (**Supplementary Fig. 3**). This observation indicated towards the monomeric state of α-Syn inside the liquid droplets at d2. In contrast, independently prepared fluorescence-5-maleimide labeled fibrils showed significantly lower lifetime values (τ_m_∼1 ns; denoted with blue colored pixels) (**Supplementary Fig. 3**), which is consistent with previously observed α-Syn fibrils in C.elegans^64^. When FLIM images of droplets at d5 and d10 were acquired; strikingly, the data showed lower τ_m_ (∼2 ns; denoted with blue colored pixels) at the center (C) of the droplets compared to the periphery (P) (τ_m_∼4 ns; denoted with green colored pixels, similar to monomeric protein). However, the IC and OC regions showed intermediate values of τ_m_ (∼2.15 and ∼3.75 ns, respectively) (**Fig 1e-f**). FLIM measurements along different Z-planes for d5 droplets further revealed a steady increase of the lifetime for planes progressively farther from the central plane, indicating that the vast majority of proteins were aggregated in the central core region of the droplet (**Supplementary Fig. 3**). During ageing (at d15 and d20), the τ_m_ of the entire droplet was, however, lowered and became close to ∼1 ns (similar to α-Syn fibril state, **Fig 1e-f**) suggesting a homogenous phase where proteins are mostly aggregated over entire droplets.

**Figure 1:**
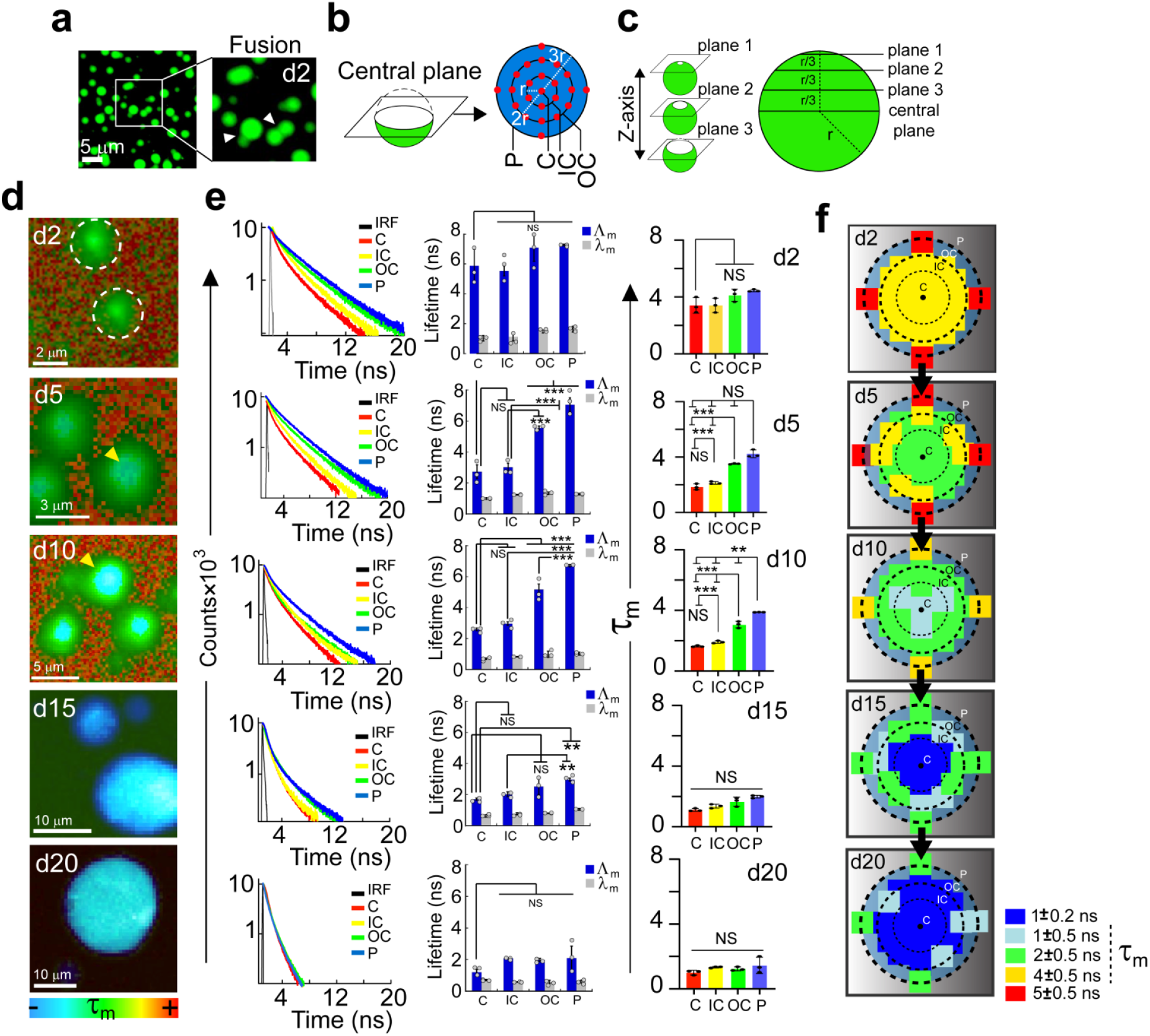
Changes in fluorescence lifetime initiates at the center and progresses towards the periphery of α-Syn phase-separated droplets during liquid-to-solid transition. **a.** Liquid droplets formed by fluorescein-5-maleimide labeled 74C-α-Syn at day 2 (d2). The inset indicates the fusion events between the droplets confirming their liquid-like state. **b.** Schematic representation of the data-points selected from the central plane of a droplet for FLIM studies. The data-points are denoted as red circles (●). **c.** Schematic showing analysis of droplets along the depth (Z-axis). Apart from the central plane, three other planes (plane 1, 2 and 3) along the Z-axis of the droplets are also imaged for FLIM to examine the distribution of lifetimes along the depth. The planes are equally spaced with a distance of *r/3* with radius of the droplet is *r*. **d.** *Left panel:* Representative color coded original FLIM images confirming the presence of the fluorescein-5-maleimide labeled 74C-α-Syn droplets at various time points (d2-d20) during liquid-to-solid transition. The color spectrum represents the range of overall mean lifetime (τ_m_) (blue being the lowest and red being the highest). The images are acquired from the central plane. The white dashed circles mark the boundary of the liquid droplets at d2. *Middle panel:* Time resolved fluorescence intensity decay obtained from different regions (C-Center, IC-Inner Circle, OC-Outer circle and P-Periphery) of the droplets at various time points (d2-d20). The black line represents instrument response function (IRF). The data-points are chosen as described in *panel b*. Representative spectra are shown. The experiment is performed three times with similar observations. *Right panel:* The long (Δ_m_) and short (λ_m_) lifetimes obtained from the intensity decay curved are plotted for C-Center, IC-Inner circle, OC-Outer circle and P-Periphery at different time-points (d2, d5, d10, d15 and d20). The data represents mean ± S.D. e. Corresponding mean lifetimes (τ_m_) obtained from intensity decay curve at different regions (C-Center, IC-Inner circle, OC-Outer circle and P-Periphery) of the phase-separated droplets are plotted for different time-points (d2, d5, d10, d15 and d20). The data represents mean ± S.D. The statistical significance (d-e) is calculated using one way ANOVA followed by Student-Newman-Keuls (SNK) post hoc test with a 95% confidence interval for n=3 independent experiments. f. Schematic representation of a lifetime map (SRM) of the droplets for different time points (d2-d20) obtained from the calculated τ_m_ values for each region (C, IC, OC, P) is shown. Each data-point is represented as a color-coded box (▪) with respect to the obtained τ_m_. Black dot (●) represents the center of the droplet and black dashed circles represent the boundary of IC, OC and P.

**Figure 2:**
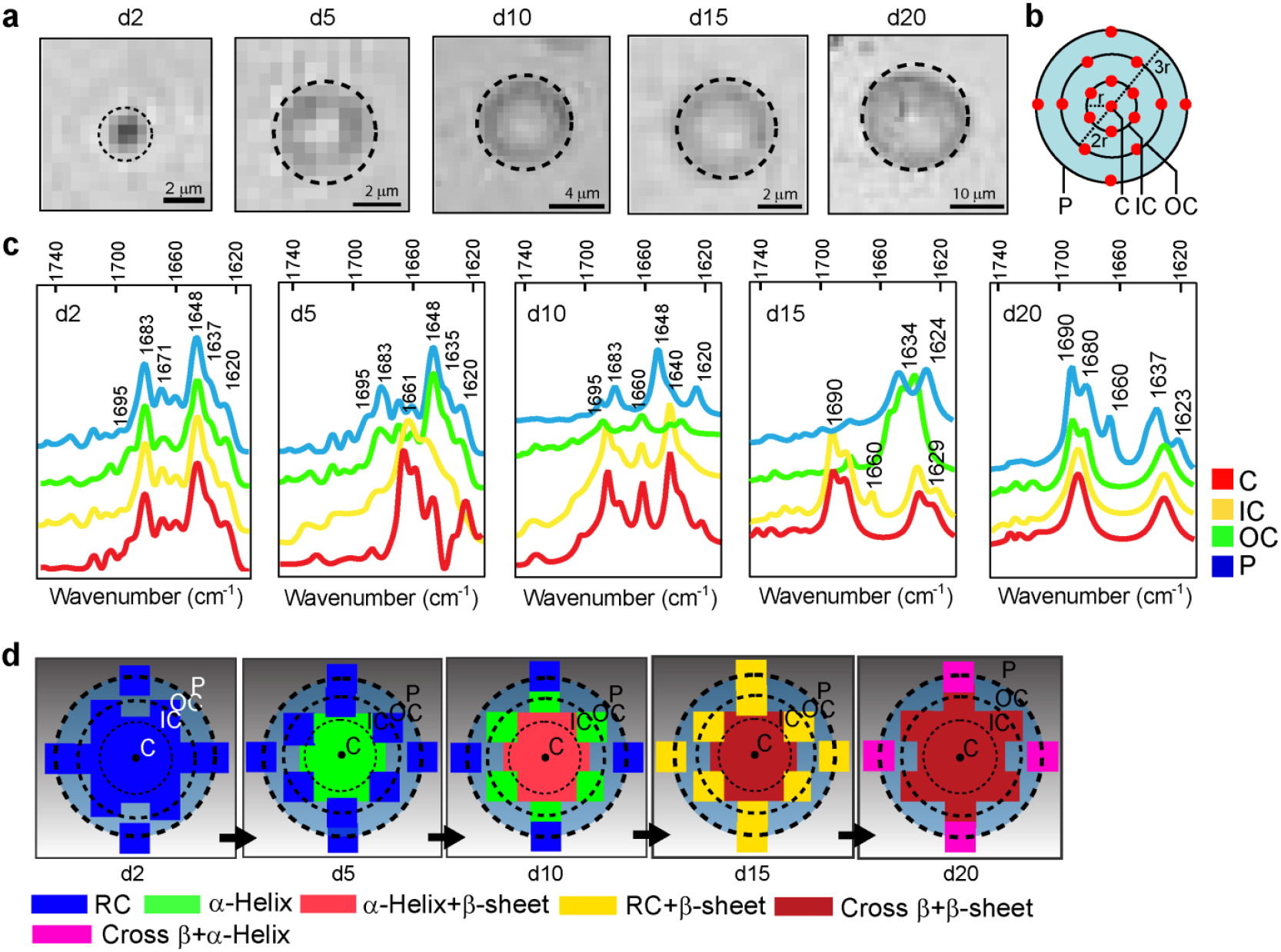
Position specific secondary structural transition in individual α-Syn phase-separated droplets during liquid-to-solid transition. **a.** FTIRM images from the central plane of phase-separated α-Syn droplets at different time-points (d2-d20) during liquid-to-solid transition. Black dashed circle represents the boundary of the droplet. The size of one single pixel was of 0.5 μm length. Representative images are shown. The experiment is performed three times with similar observations. **b.** Schematic representation of the data-points selected from the central plane of a droplet for FTIRM studies. The data-points are denoted as red dots (●). **c.** Corresponding FTIR spectra obtained from different regions (C-Center, IC-Inner Circle, OC-Outer circle and P-Periphery) of the droplets are plotted for each time point (d2-d20). One representative FTIR spectra for each data-point is shown. The respective wavenumber (cm^-1^) for every peak in each spectrum is denoted. The experiment is carried out three times with similar results. **d.** Schematic representation depicting a spatially resolved secondary structural (based on deconvolution and % secondary structure calculation) map (SRM) of the droplets for different time during incubation (d2-d20). Each data-point, from different regions, is represented by a color-coded box (▪) with respect to the obtained % secondary structural majority. Black dot (●) represents the center and black dashed circles represent the boundary of IC, OC and P.

**Figure 3:**
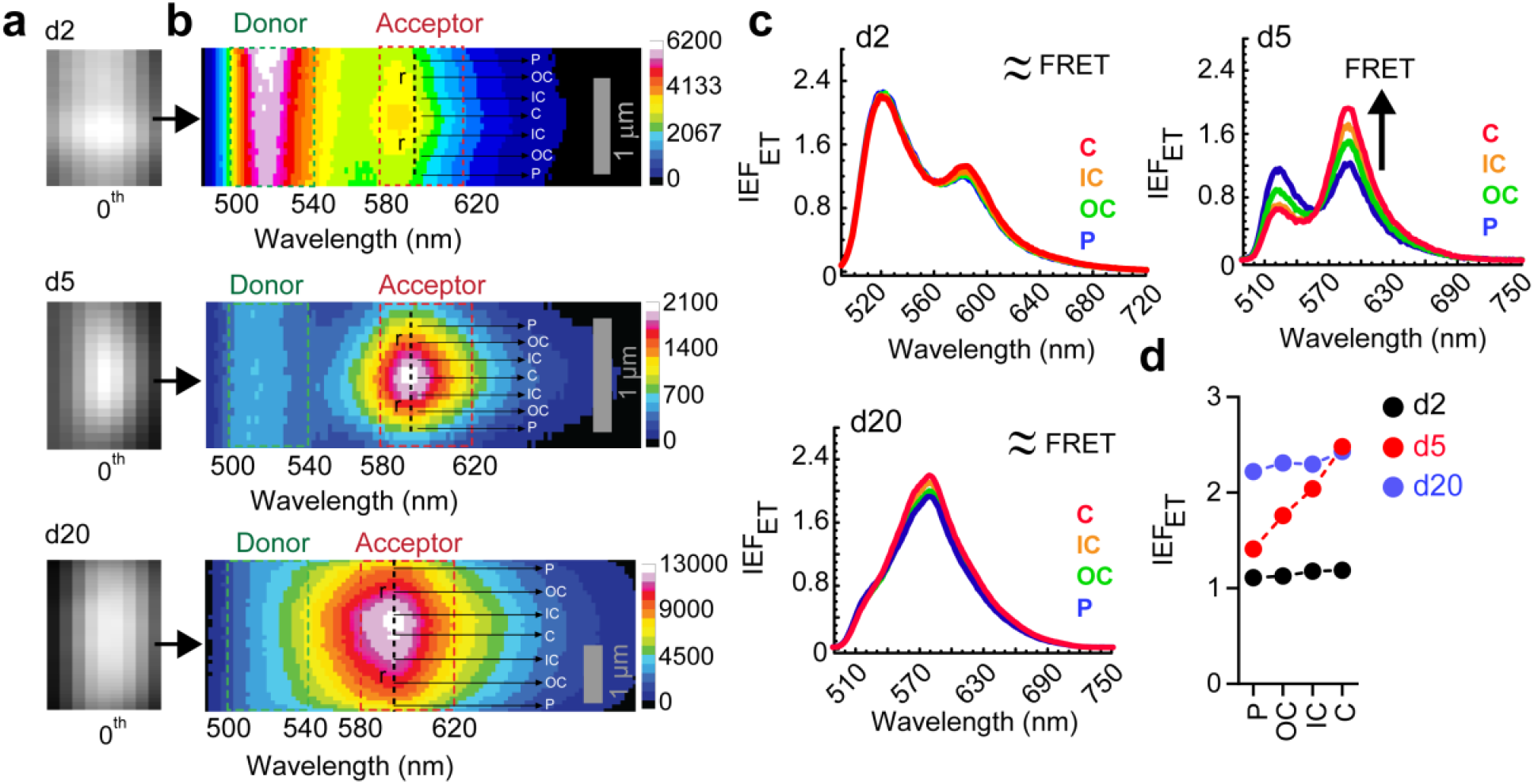
Intermolecular interaction responsible for protein aggregation initiates at the center of α-Syn phase-separated droplets during liquid-to-solid transition. **a.** The 0^th^ order images of α-Syn liquid droplets at various time points (d2, d5 and d20) during liquid-to-solid transition are shown. From these images, the spectrally resolved images are obtained. **b.** Spectrally resolved microscopic images of α-Syn liquid droplets at d2, d5 and d20, containing both fluorescein-5-maleimide (donor, *D*) and rhodamine-C2-maleimide (acceptor, *A*) labeled 74C-α-Syn. The X-axis represents wavelength (nm) and Y-axis represents distance (μm). The color codes of the pixels represent emission fluorescence intensity values in arbitrary units (a. u.). The *D* and *A* fluorescence emission regions are marked with green and red boxes, respectively. The radii of the droplets on the spectral images are marked with black dashed lines (---). **c.** The *IEF_ET_* values are plotted against the emission wavelengths for different regions (C, IC, OC and P) for d2, d5 and d20 droplets. **d.** The maximum *IEF_ET_* values (measure of FRET) are plotted for P, OC, IC and C for d2, d5 and d20. The experiment is carried out 2 independent times with similar results.

We found similar results showing the formation of a solid-like core with reduced lifetime at d5, which gradually propagated towards the periphery till d20 when we performed FLIM experiments with sample prepared in 2 ml Eppendorf tubes. Further, protein concentration also remained same after 20 days incubation in microscopy slides due to incubation in moist chamber and the FLIM results was not largely affected by sedimentation/wetting of the droplets on glass surface (we used only spherical droplet for the measurements) (**Supplementary Fig. 4-6**). Some heavier droplets at d10 onwards did show interaction with the surface at the bottom of the slide and showed a distorted (ring-like) lifetime distribution (**Supplementary Fig. 6**).

Collectively, our data showed that the local microenvironment of α-Syn was homogeneous throughout the droplets at early stages of LLPS with α-Syn of higher τ_m_ (**Fig 1f**). However, it began to change at the center of the droplets first and at later time-points, entire droplets became homogeneous in terms of reduced τ_m_ values (**Fig 1f**) similar to fibrils.

### Conformational transition of α-Syn starts at the center and propagates towards the periphery of the phase-separated droplets

Subsequently, to probe the secondary structure of α-Syn at different regions in the droplets during liquid-to-solid transition, we employed Fourier transform infra-red microscopy (FTIRM) imaging in a single-point mode (**Supplementary methods**). This was done to probe whether the dynamic progression of the altered local microenvironment (**Fig 1**) is due to conformational transition of α-Syn associated with aggregation. Unlabeled wild-type (WT) α-Syn liquid droplets were imaged using FTIRM at various time-points (d2-d20) **(Fig 2a)**. The data-points were chosen at C, IC, OC and P of the microscopic image of the central plane of the droplet and corresponding FTIR spectra were recorded **(Fig 2b-c** and **Supplementary Fig. 7**). The obtained spectra were deconvoluted and the individual peaks were assigned for specific secondary structures and their abundance (**Supplementary Fig. 7-8**). To make sure deconvolution does not affect the signal to noise ratio, water subtraction from each of our spectra by taking the appropriate buffer as a control was performed. In our case, we did not find any major peaks corresponding to artifacts or water vapor in the region beyond 1700 cm^-1^ (**Fig 2c**). Some small peaks were observed which may be due to the side chain functional groups from acidic amino acids and carbonyls.

Our result revealed identical signature for all regions (C, IC, OC, P) with the highest intensity peak at 1648 cm^-1^, corresponding to the random coil (RC) conformation in α-Syn droplet at d2 (**Fig 2c and Supplementary Fig. 7-8**). The individual peaks of the deconvoluted spectra further confirmed that the FTIR signature resembled soluble, monomeric α-Syn (**Supplementary Fig. 7**). Interestingly, unlike the early stages of LLPS (d2); at d5, the regions OC and P showed a clear difference in their respective FTIR spectra compared to C and IC (**Fig 2c** and **Supplementary Fig. 7**), which was consistent with the lifetime data (**Fig 1**). While OC and P showed mostly RC conformation, C and IC showed presence of α-helical conformation (**Fig 2c and Supplementary Fig. 7-8**). In line with previous reports, this suggested that α-Syn LLPS and subsequent aggregation is mediated by α-helical oligomeric intermediate^55, 65^. At d10; C, IC and OC exhibited β-sheet and α-helical structures as major conformations (**Fig 2c** and **Supplementary Fig. 7-8**). The periphery (P) of the droplets at d10 still showed majorly RC conformation (**Fig 2c** and **Supplementary Fig. 7-8**). At d15, FTIR spectra from C and IC had two major peaks corresponding to β-sheet and cross β-sheet structure (**Fig 2c** and **Supplementary Fig. 7-8**). OC showed both RC and β-sheet; while P showed β-sheet and RC conformations **(Fig 2c** and **Supplementary Fig. 7-8)**. The d10 and d15 data clearly indicated that structural transition leading to aggregation was initiated at the center. At d20, the entire droplets were found to exhibit only β-sheet and cross β-sheet structures (**Fig 2c** and **Supplementary Fig. 7-8**) indicating the progression of aggregation towards the periphery consistent with our FLIM data. The sudden increase in the % abundance of α-helical conformation (**Supplementary Fig. 8**) at the periphery (P) at d20 might be due to the recruitment of fresh molecules and their further conversion into α-helix rich state after interacting with β-sheet rich droplets. The % abundance of the secondary structures and FTIR peak assignments at different spatial locations of the droplets during liquid-to-solid transition are provided in **Supplementary table 2** and **3**. Notably, a very similar method using Raman microscopy has been recently employed to calculate the absolute concentrations of ataxin-3 protein inside a single droplet^66^ further supporting the robustness of our FTIRM experiments.

In essence, our spatially-resolved FTIRM measurements on single droplets suggested that α-Syn liquid-to-solid transition was accompanied by conformational transition (RC◊α-helix◊β-sheet), which initiates at the center and then gradually covers the entire droplet (**Fig 2d**).

### Single droplet FRET showed stronger intermolecular interaction at the center promoting liquid to solid transition α-Syn liquid droplets

Next, to further probe that the solid-like core formation at the center was due to α-Syn aggregation, we performed spatially as well as spectrally resolved FRET^55^ for single α-Syn liquid droplets during liquid-to-solid transition (**Supplementary Fig. 9**). We chose fluorescein-5-maleimide labeled (10% v/v) 74C-α-Syn as donor (*D*) and rhodamine-C2-maleimide labeled (10% v/v) 74C-α-Syn as acceptor (*A*) for our microscopy based FRET study at single droplet resolution^55^. The 74^th^ position was chosen to delineate intermolecular interactions of the NAC domain responsible for protein aggregation^55, 63^ in a location specific manner inside the droplets. Crowding (10% w/v PEG-8000) induced α-Syn droplets were generated using an equimolar (200 μM) concentration of *D* and *A*. The spectral images (**Supplementary methods**) of liquid droplets containing both *D* and *A* were analyzed at C, OC, IC and P regions (**Fig 3a-b)**. At d2, we observed similar extent of apparent FRET efficiency^55^ as evident from the *IEF_ET_* values (∼1.2) at different locations (C, IC, OC, P) indicating the similar extent of intermolecular interaction throughout the droplets at early stages of LLPS (**Fig 3c-d**). Strikingly, at d5, we found that the apparent FRET efficiency (*IEF_ET_*) was highest at the center (C, *IEF_ET_* ≈ 2.4) and progressively lower towards the periphery (P, *IEF_ET_* ≈ 1.4) (**Fig 3c-d)**. This clearly indicated that the solid-like core at the center forms due to inter-domain interaction of the NAC region, which eventually results in amyloid aggregation. At d20, all the regions (C, IC, OC and P) showed similar *IEF_ET_* values (*IEF_ET_* ≈ 2.4) (**Fig 3c-d)** however with the overall increased magnitude of the *IEF_ET_* **(Fig 3d)**. The data clearly indicates that higher extent of intermolecular interaction at the center leads to liquid to solid-transition, which eventually populates the entire droplet.

### Spatiotemporal FRAP shows diffusivity of α-Syn molecules are slower at the center compared to the periphery of the droplets

To find more evidence supporting the presence of a solid-like core, we also performed FRAP at different locations (C and OC) inside the same α-Syn droplet at different time-points (d2, d5, d7 and d15) (**Fig 4a**). Furthermore, to provide additional evidence for the formation and maturation of the solid-like core, we have now performed spatiotemporal FRAP with NHS-rhodamine labeled (10% v/v labeled) α-Syn droplets. Briefly, 200 μM of α-Syn was phase separated in presence of 10% (w/v) PEG-8000 in 2 ml eppendorf tubes (as mentioned in the manuscript). At regular time intervals (d2, d5, d7 and d15), 5 μl LLPS sample was drop-casted on a clean glass-slide and subjected to FRAP measurements. The droplets were bleached at three regions along their diameter: R1- (center-C), R2- (away from center-OC) and R3- (at the periphery-P) (**Fig 4a**). Since the d2 droplets were very small, due to the resolution limit of the confocal microscope, no spatially resolved bleaching could be performed (**Fig 4b-c**).

**Figure 4:**
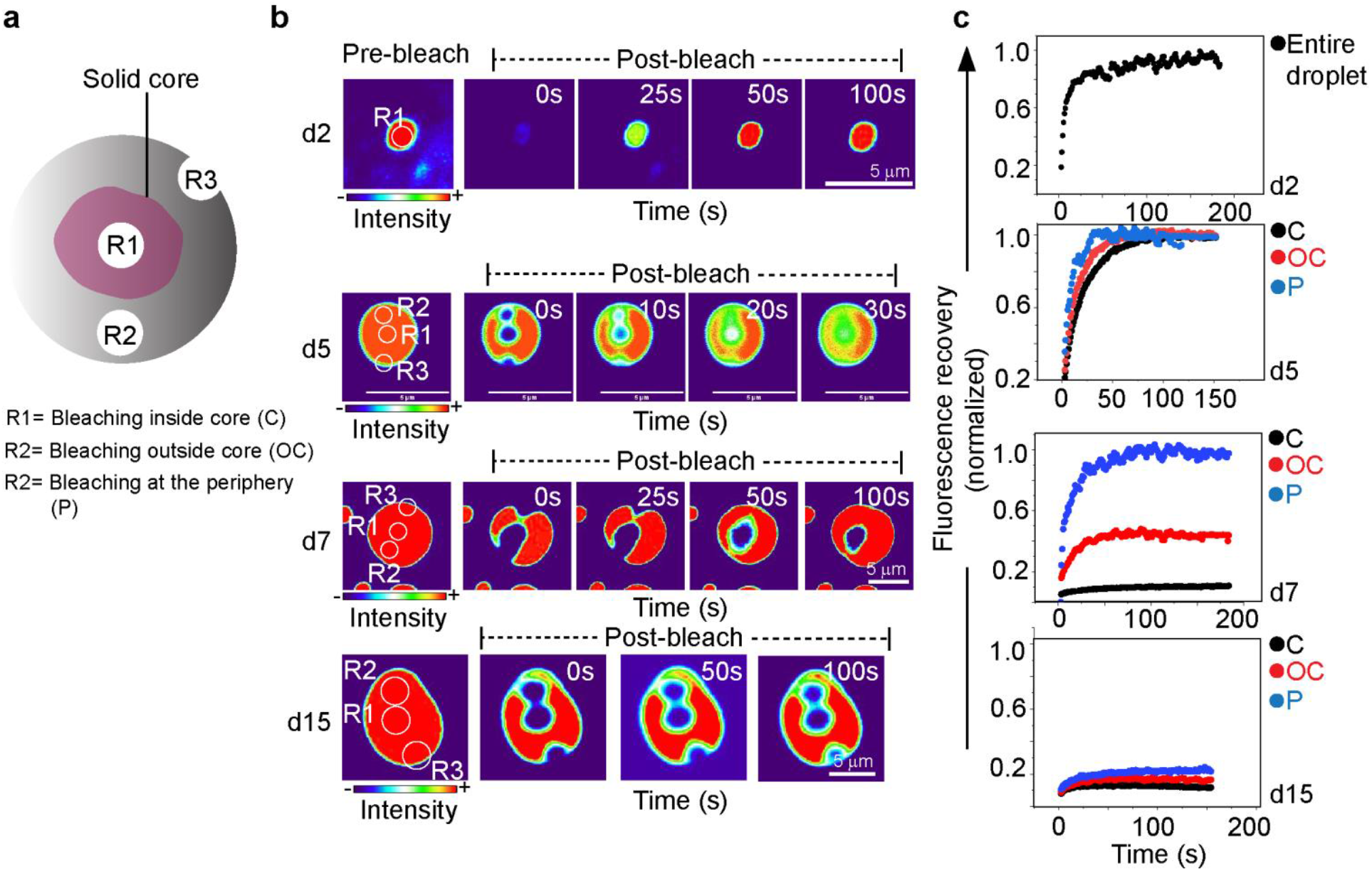
FRAP at different locations of α-Syn phase separated droplet during liquid-to-solid transition. **a.** Schematic showing the bleaching regions chosen for our FRAP study. R1 represents a bleaching region at the center of the droplets where the solid-like core is present. R2 represents a bleaching region away from the core (at the OC region) where the molecules are supposed to be relatively more diffusive. R3 is at the periphery where the molecules are most diffusive. **b.** Representative time-lapse images of the droplets at d2, d5, d7 and d15 during FRAP are shown. The droplets are bleached at C (R1), OC (R2) and P (R3) regions which are marked with white circles. Notably, d2 droplets were small and could not be bleached in a spatially resolved manner. The fluorescence recovery is shown in thermal LUT for better visualization of the apparent differences in the rate of the fluorescence recovery for different regions. **c.** The normalized fluorescence recovery is shown for C, OC and P regions at different time-points (d2-d15). The normalized recovery of the entire droplet is shown for d2.

At d5, d7 and d15, the droplets become fairly large to perform FRAP at different locations along the diameter without any significantly overlapping bleaching region. Our results showed that at d2 and d5, all the regions (C, OC and P) showed complete fluorescence recovery. However, the kinetics/rate of fluorescence recovery was considerably slower at the center (C) followed by OC and P regions for d5 droplets (**Fig 4b-c**). The fluorescence images were converted to thermal LUT to probe the difference in the fluorescence recovery at the C region compared to the OC and P regions (**Figure 4b-c** and **Supplementary video 1**). At d7, the fluorescence intensity did not recover when the droplets were bleached at the C region (**Fig 4b-c**). Strikingly, the OC region of the same droplet showed almost 50% recovery of fluorescence indicating that the molecules away from the C region were relatively more diffusive. At the P region, the recovery was almost ∼100% (**Fig 4b-c**). Our data clearly points out that at the C region, because of aggregation; the α-Syn molecules were restricted compared to the molecules at the periphery of the droplets at d5 and d10 which was very consistent with our FLIM, FTIR, FRET observations. As expected, at d15, the C, OC and P regions showed minimal recovery of fluorescence indicating propagation of the solid-like core to the surface of the droplet (**Fig 4b-c**).

### Direct observation of solidification at the center of liquid droplets using transmission electron microscopy (TEM) imaging

To directly confirm the presence of solid-like core at the center of the liquid droplets, we performed time-dependent negative stained TEM imaging of α-Syn liquid droplets during liquid-to-solid transition (d2-d15). The grayscale intensity profiles from TEM images were analyzed using ImageJ (NIH). The detailed description of the analysis is given in the supplementary information.

At the beginning of the droplet formation (d2), TEM imaging showed circular droplet-like structures, which mostly appeared to have amorphous protein content (**Fig 5a-b**). Image analysis and subsequent grayscale intensity profiles were plotted along the diameter of the droplets, which suggested homogeneous appearance of droplet without any protein rich, electron transparent region inside the d2 droplets (**Fig 5c**).

**Figure 5:**
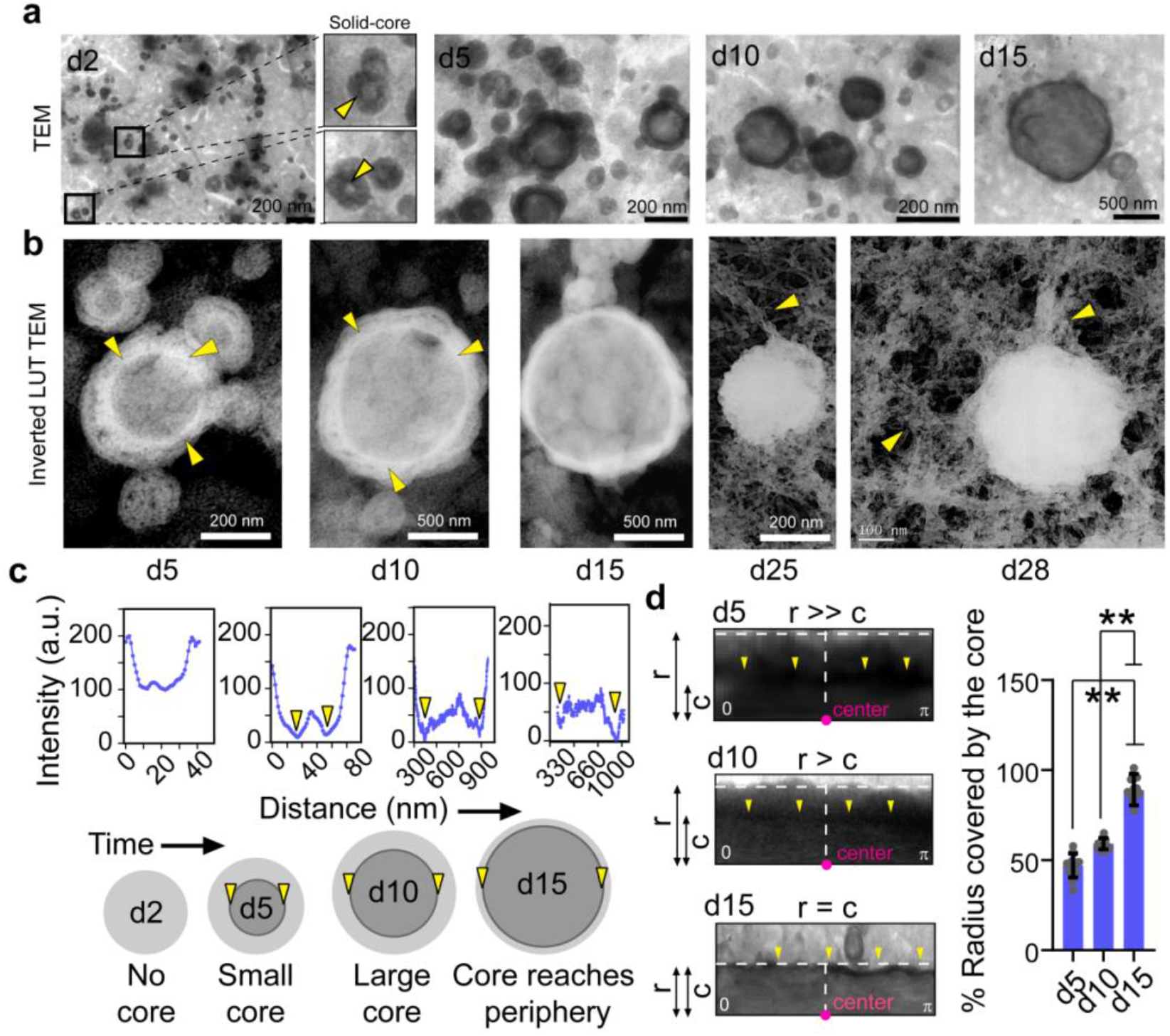
Transmission electron microscope (TEM) images of α-Syn liquid droplets during liquid-to-solid transition. **a.** TEM images of α-Syn phase separated droplets at d2 shows very few droplets which possess the core-like structures (marked with yellow triangular pointers). However, at d5 and d10, the core structure is clearly visible for most droplets. After d15, the core structure is no longer visualized. This could be due to the fact that the core reaches the periphery of the droplet with time. **b.** Representative TEM images of the droplets at d5, d10 and d15 with inverted lookup table (LUT) clearly indicates the presence of a progressively growing core inside the droplets (marked with yellow triangular pointers). At d25 and d30, fibril-like morphology emerges from the droplets (marked with yellow triangular pointers) indicating amyloid fibril formation starts inside the droplets^55^. **c. *Upper panel:*** The pixel intensity is calculated from the cross section (white dashed line) of the droplet at various time-points from d2 to d15. The yellow triangular pointers denote the intensity drops indicating the start and end of the boundary of the solid-like core structure. ***Lower panel:*** Schematic representation of the droplets at different time-points (d2-d15) where the d2 droplet showing the absence of the solid-like core. The core structure appears at d5 and extends towards the periphery with subsequent incubation. The yellow triangular pointers denote the boundary of the core at each time-point where the intensity drop is observed. **d. *Left panel:*** Angular maps (0◊180°) of the droplets at d5, d10 and d15 from the center (●) to the periphery (white dashed horizontal line) are generated. The yellow triangular pointers denote the boundary of the core at each time-point. The radius of the entire droplet is denoted as “r” and the radius of the core is denoted as “c”. Visual observations suggest that at d5, r>>c; at d10, r>c and at d15, r ≈ c. ***Right panel:*** The % radius of the droplet that is covered by the solid core showing significantly higher core at d15 (∼90%) compared to d10 (∼60%) and d5 (∼50%). The values represent mean ± S.D. The statistical significance is calculated using two tailed paired Student-t-test with a 95% confidence interval for n=20 individual droplets. The experiment is performed three times with similar results.

Interestingly, the TEM images of d5 droplets clearly showed formation of a protein rich structure (electron transparent region in negative staining) at the center with a clear ring-like boundary inside the droplets (**Fig 5a-b**). This could be the solid-like core that formed at the center of the droplets after d5. The signal intensity profiles along the diameter of the droplets showed sharp grayscale intensity drops corresponding to the boundary of the core structure (**Fig 5c**). The solid-like core structures further grew in size when droplets were allowed to age (**Fig 5a-b**). Eventually, after d25, amyloid fibrils started emerging from the droplets as evident from our TEM imaging (**Fig 5b**). Our observation was further evident from the distance between the two grayscale intensity drops in **Fig 5c**. We further quantified the % of radius of the solid core based on the electron transparent area at the center of the droplets during time (d5, d10 and d15) using images of multiple droplets (**Fig 5c**).

We converted the TEM images into angular maps (0-180°) from the periphery to the center to ease the quantification (**Fig 5d, *Left panel*** and **Supplementary Fig. 10)**. This operation straightened the images for easy visualization and quantification of the radial average intensity profile of a curved object^67^. The angular maps quantification data indicated that at d5, the solid-like core covered ∼50% of the radius. At d10, the core progressed and covered up to ∼60% of the droplet radius and finally, at d15, the aggregate core covered almost the entire droplet (∼90% of the radius) (**Fig 5d, *Right panel*)**.

Our TEM data thus conclusively proves that the initiation and subsequent progression of α-Syn aggregation inside the droplets starts at the center and reaches the periphery during liquid-to-solid transition of α-Syn LLPS. Additionally, cryo-SEM imaging of d5 droplets also revealed a core-like structure at the center of the α-Syn droplets (**Supplementary Fig. 11**) further supporting our TEM observations.

### Fusion of different sized droplets contributing towards droplet growth and liquid-to-solid transition

Furthermore, we were interested to know how droplet fusion could contribute to the growth and subsequent solidification of α-Syn LLPS. We hypothesized that the larger droplets might acquire solid-like core faster than the smaller droplets. Subsequently during fusion, the smaller droplets without the structured core could contribute their contents for further growth of larger droplets (due to difference in their thermodynamic stability). To examine the possible differences in the structured core, we carefully chosen the fusion events of droplets formed by fluorescein-5-maleimide labeled 74C-α-Syn (10% v/v) where one droplet was larger compared to the other droplet undergoing fusion at d5 (LD1-larger droplet, LD2-smaller droplet). We performed FLIM at selected data-points (p1-p6) along the longitudinal fusion axis, which connects the centers of the droplets (p3 and p6) with the fusion region (p1) (**Fig 6a, *Left panel***). The microscopic observation of the droplets using FLIM showed distinct differences in the τ_m_ at different regions of the droplets (**Fig 6a, *Right panel***). Further, the time-resolved fluorescence intensity decay from each of the data-points were collected and fitted with bi-exponential decay function (**Fig 6b, *Left panel***). The decay profiles and corresponding τ_m_ data suggested that the central region of both the droplets had lower τ_m_ (p6 ∼1.3 ns and p3 ∼2 ns) while the periphery had higher τ_m_ (∼3 ns). Interestingly, the fusion region showed similar τ_m_ as the periphery (∼3 ns) (**Fig 6b, *Right panel***). In addition, the center of the large droplet showed much less τ_m_ compared to the smaller droplet suggesting that the larger droplet possessed a more stable structured core for solidification compared to the smaller droplets. This is further consistent with FTIRM data of two different sized droplets undergoing fusion at d5 (**Fig 6c**).

**Figure 6:**
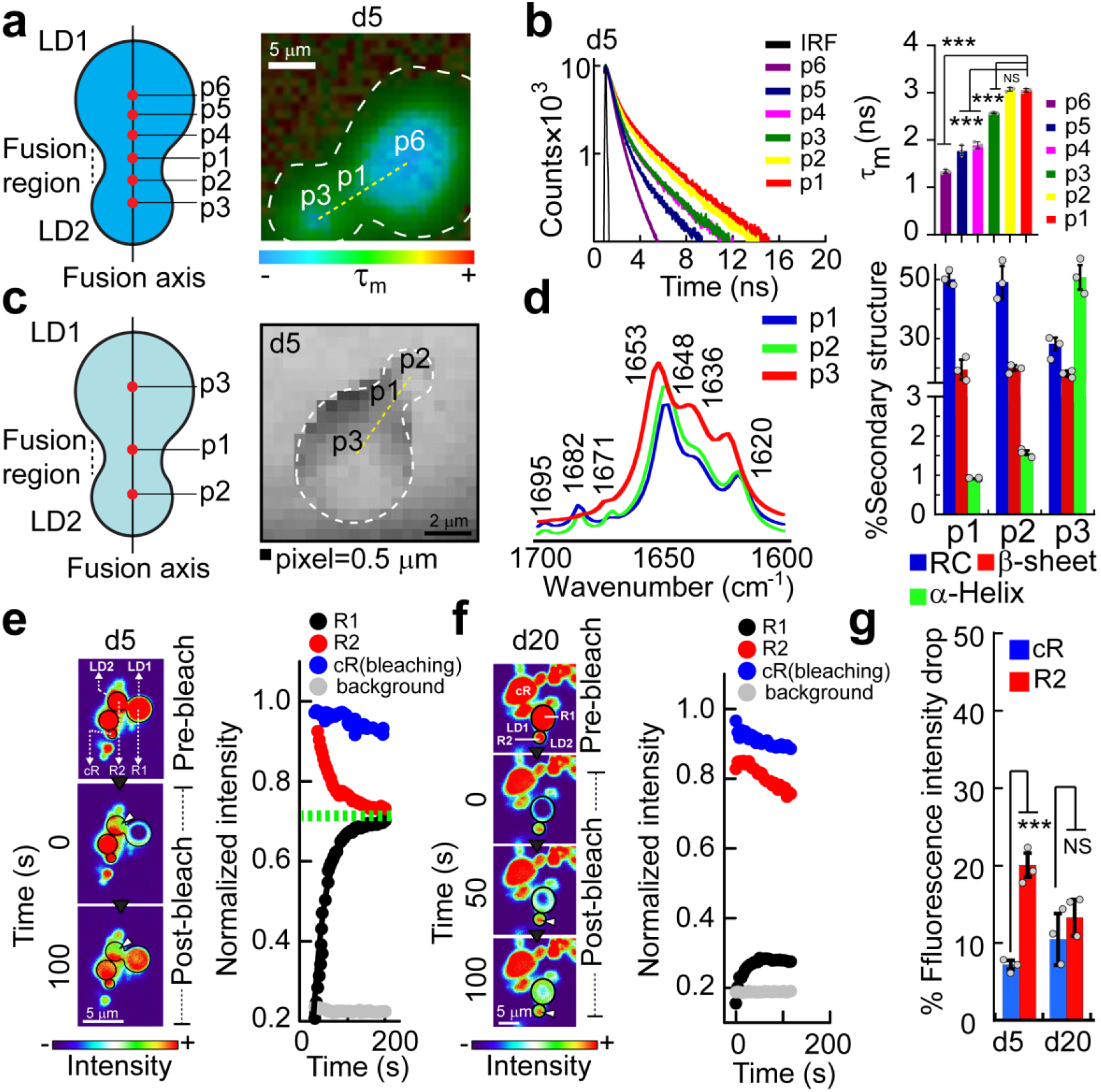
Growth of the larger droplets containing structured core at the center upon fusion with smaller droplets. **a.** Schematic (***left panel***) and corresponding original FLIM image (***right***) showing a fusion event of two heterogeneous sized droplets at d5. The boundary of the fusion event and longitudinal fusion axis (***right panel***) is marked with a white dashed curve and yellow dashed line, respectively. The color spectrum represents the range of overall mean lifetime (τ_m_) (blue is the lowest and red is the highest). **b. *Left panel:*** The time resolved fluorescence intensity decay profiles obtained from different regions (from p1 to p6) on the fusing droplets. The black line represents the IRF. Different color codes are assigned to denote different positions from p1 to p6. ***Right panel:*** The τ_m_ for each data point (from p1 to p6) is plotted. Values represents mean ± S.D for n=3 independent experiments. The significance is computed using one way ANOVA followed by Student-Newman-Keuls (SNK) post hoc test with a 95% confidence interval for n=3 independent experiments. **c.** *Schematic* (***left panel***) and single point FTIRM (***right panel***) image of fusion event between two heterogeneous sized droplets (liquid droplet 1 (LD1) and liquid droplet 2 (LD2)) at d5. The boundary of the fusion event and longitudinal fusion axis (right) is marked with a white dashed curve and yellow dashed line, respectively. **d. *Left panel:*** Corresponding FTIR spectra obtained from different regions (p1-p3) of the fusion event at d5. The respective wavenumber (cm^-1^) for every peak in each spectrum is denoted. Representative spectra are shown. The experiment is carried out three times with similar results. ***Right panel:*** The % abundance of the secondary structures (RC, α-helix and β-sheet) for each of the data-points (p1, p2 and p3) is plotted. Values represent mean ± S.D for n=3 independent experiments. **e-f. *Left panel*:** Representative images of FRAP by two fusing droplets (LD1 and LD2) at d5 and d20. Liquid droplet 1 (large, LD1) is actively bleached at region ‘R1’ (diameter of bleaching region = 3 μm). The fluorescence intensity recovery post bleaching is acquired from both R1 in LD1 and R2 in LD2. The gradual intensity decrease due to prolonged laser exposure (passive bleaching) is quantified from an independent region of interest (cR) on a different droplet. The images are represented in “thermal” pseudo-color. The color spectrum represents fluorescence intensity (blue, lowest and red, highest). **e-f. *Right panel*:** Corresponding post-bleach fluorescence intensities of R1, R2, cR and background. At d5 (**e**), the post-bleach intensity of R1 gradually increases with time (recovery) while the intensity of R2 gradually decreases. The intensity of R1 and R2 eventually reaches equilibrium (denoted as green dashed line). At d20 (**f**), the post-bleach intensity of R1 neither recover with time nor the intensity of R2 substantially decreases (only decreases to some extent due to passive laser bleaching). Representative data is shown. The experiment is performed three times with similar results. **g.** The % decrease in the fluorescence intensity of cR (due to passive laser bleaching) and R2 (due to molecule transfer to the neighboring fusion partner) are plotted at different time-points (d5 and d20). The data showing the % fluorescence intensity drop of R2 is significantly higher than passive laser bleaching (cR) at early time-points (d5). The values represent mean ± S.D. The statistical significance is calculated using two tailed paired Student-t-test with a 95% confidence interval for n=3 independent experiments.

The FTIR spectra from the images of two fusing droplets at d5 showed majorly RC conformation for p1 (at the fusion junction) and p2 (center of smaller droplet). The center of larger sized droplets, p3 however showed α-helix conformation (**Fig 6d, *Left panel*** and **Supplementary table 2**). The % abundance of the secondary structure at different regions is provided in **Fig 6d, *Right panel*** and **Supplementary table 4**. This observation suggests that immature (smaller, thermodynamically unstable) droplets tend to fuse with more stable (bigger) droplets containing a stable solid-like core.

To probe that smaller droplets without structured core supply the content to larger droplets for their further growth, we performed FRAP of droplets undergoing fusion events at d5. The early stage of droplet fusion events was chosen to ease detection of substantial differences (if any) between two fusing droplets (**Fig 6e, *Left panel***). We used NHS-rhodamine labeled (10% labeled; v/v) α-Syn as passive photobleaching of rhodamine is much less compared to fluorescein^68^. The larger droplet undergoing fusion was actively bleached with laser and its fluorescence recovery was observed. Strikingly, we noticed, as the bleached droplet recovered, the intensity of its unbleached partner (smaller droplet) was substantially decreased. Moreover, the fluorescence intensities of both the droplets eventually reached equilibrium (**Fig 6e, *Right panel***). For example, at d5, the intensity of the actively bleached, larger droplet was recovered up to ∼80% but simultaneously, the intensity of the unbleached, smaller partner droplet was decreased by ∼20%. The passive bleaching effect by the monitoring laser was negligible (∼8%). This could be because the un-bleached droplet was constantly providing the new α-Syn molecules to bleached droplet. Overall, the data provides dynamic, real time evidence that inter-droplet transfer of α-Syn occurs during fusion events at d5 and the larger droplets containing structured core indeed grow and solidify during time with expense of small droplets after fusion. In contrast, when similar FRAP experiments were performed with neighboring droplets at d20, no substantial fluorescence recovery of the larger droplet was observed. The unbleached, smaller droplet in proximity also did not show any decrease in fluorescence intensity (except due to passive bleaching by the monitoring laser) (**Fig 6f-g**). Our observations suggest that at later stages of LLPS, the rigidity of the droplet increases in expense of the molecular diffusion, and as a result, the mass transfer between two droplets in proximity is also stopped. Together, the data shows that although α-Syn aggregation initiates at the center of the droplet, the monomeric, soluble-like α-Syn participates in the molecular exchange during early stages of fusion.

## Discussion

Previously, many proteins undergoing LLPS showed the phenomenon of liquid-to-solid transition during time^12, 13, 42–44, 46–49^. The solidification of liquid droplets not only aids normal biological functions^47–49^ of many liquid droplets but also promote toxic protein aggregation associated with various neurological disorders as shown for FUS^41, 42^, Tau^43, 44^, TDP-43^57^ and α-Syn^55^. In the phase-separated droplets undergoing solidification, all molecules might not be in exactly similar microenvironment. Also, it is unlikely that liquid-to-solid transition of protein LLPS is a simple two-state process, where all protein molecules in liquid state are transitioned into solid state at once. Rather, the pathway of liquid-to-solid transition might resemble crystallization process where at first; a nucleus is formed, which gradually recruits molecules for further crystallization^69^. We asked, “Where exactly does solidification (nucleation event) begin for liquid droplets undergoing liquid-to-solid transition?”

Our experiments on α-Syn droplets showed that at the beginning of droplet formation, all α-Syn molecules possess similar microenvironment and have monomeric (RC) conformation (**Fig 1-2**). During maturation of α-Syn LLPS, the local microenvironment begins to change at the center (with reduced lifetime of fluorophore attached to α-Syn). This change could be associated with α-Syn assembly as evident from decrease in fluorescence lifetime and conformational transitions. Previous reports have shown that molecular entanglement during α-Syn aggregation results in self-quenching^70^ of the attached fluorescent probe due to close proximity^64^. This leads to a reduced fluorescence lifetime during aggregation and fibril formation. During time, the center of the droplet shows conformational transition into α-helix-rich state, an intermediate known for α-Syn aggregation, LLPS and gel formation^55, 65^. The local microenvironment of α-Syn at the center of the droplet further changes (reduced lifetime), which expands and covers the entire droplet with time (**Fig 1**). FTIRM imaging also shows structural transition of α-Syn favorable for aggregation and amyloid formation initiating at the center and then gradually progressing towards the periphery till entire droplet shows β-sheet-rich amyloid fibril structure (**Fig 2**). The spatially resolved, position specific measurements, therefore, enables us to identify the solid-like core at the center, which acts as a nucleation site for subsequent solidification/aggregation inside the droplets. Important to note, the kinetics of α-Syn LLPS^71^ might influence the observation of the solid-like core. Conditions that triggers immediate LLPS and accelerated liquid-to-solid transition^59, 60^ could result in formation and subsequent propagation of the solid-like core within minutes to hours. On the other hand, our method is useful to delineate the spatiotemporal changes with higher resolution because of the slow kinetics of the system.

The probable reason behind solidification starting at the center could be because the environment of α-Syn molecules at the surface of the droplet being very different from that of the interior. Each α-Syn molecule at the interior of the droplet is surrounded by many other α-Syn molecules in all directions, resulting in very less amount of net force exerted on these molecules residing deep in the center of a droplet compared to the surface. In contrast, α-Syn molecules at the surface of a droplet have asymmetrical environment due to close proximity of the boundary where they experience strong interaction on one side (towards the center) and minimal/no interaction on the other side (outside). Moreover, because the phase-separated bodies are membraneless, there could also be very fast exchange of the surface α-Syn molecules with the surrounding. High surface energy of such α-Syn molecules makes them energetically unstable and therefore, prevents stable assembly. Moreover, increased local concentration of α-Syn at the center might also trigger self-assembly faster than the other places (such as periphery) of the droplet. This is shown in concentration distribution in PGL3 condensates^72^ and LLPS associated with functions such as heterochromatin assembly^49^—indicating that the presented phenomenon could be the generic property for many protein condensates undergoing liquid-to-solid transition.

Indeed, our FRET analysis suggests that the intermolecular NAC-domain interaction responsible for amyloid aggregation plays a crucial role in forming the solid-like core, which initiates at the center on d5 and extends towards the surface of the droplet during liquid-to-solid transition (**Fig 3**). This is further supported by the difference in the % recovery of fluorescence at different location of the same droplet (**Fig 4**). Indeed a core-like structure is also observed under TEM at earlier time-points (**Fig 5**). However, at this point, we cannot exclude the possibility of multiple nucleation (but very small in size) events, which are close to each other that, may occur inside the droplet simultaneously.

During droplet maturation, three distinct processes generally occur: continuous consumption of new α-Syn molecules from the solution into the droplet; Ostwald ripening^73^ and droplet fusion^74^. Smaller and thermodynamically unstable droplets tend to fuse with larger, stable droplets to minimize free energy^4^. We hypothesize that during fusion, the transfer of constituent molecules takes place from smaller to larger droplets containing solid-like, aggregate core structure.

Using FLIM and single point FTIRM, we show that indeed, the larger droplets possess a solid-like core (low lifetime and more structured α-Syn) at the center, which the smaller droplets lack. This indicates a heterogeneous nature of α-Syn LLPS (**Fig 6**). Smaller, homogeneous droplets fuse with larger droplets where the aggregation process has already been started. This means that at any given time, the droplets, which contain a solid-like core contribute towards the progression of liquid-to-solid transition at the cost of the smaller, soluble droplets, which is consistent with our FRAP study. This phenomenon is however completely hindered at the end of solidification because the aggregation of α-Syn reaches the periphery of the droplets preventing further molecular diffusion (**Fig 6**). This observation is in line with previous reports showing an arrested, supersaturated state of protein droplets where the growth of the droplets is completely stopped^6^.

The present data suggests that solidification/aggregation of α-Syn liquid droplets initiates at the center due to facilitated self-assembly and/or conformational transition (**Fig 7**) in a spatiotemporal manner. The current work might help to understand liquid-to-solid transition and fibril formation by many other proteins containing IDRs, which are associated with neurological disorder as well as functional membraneless organelles formation.

**Figure 7:**
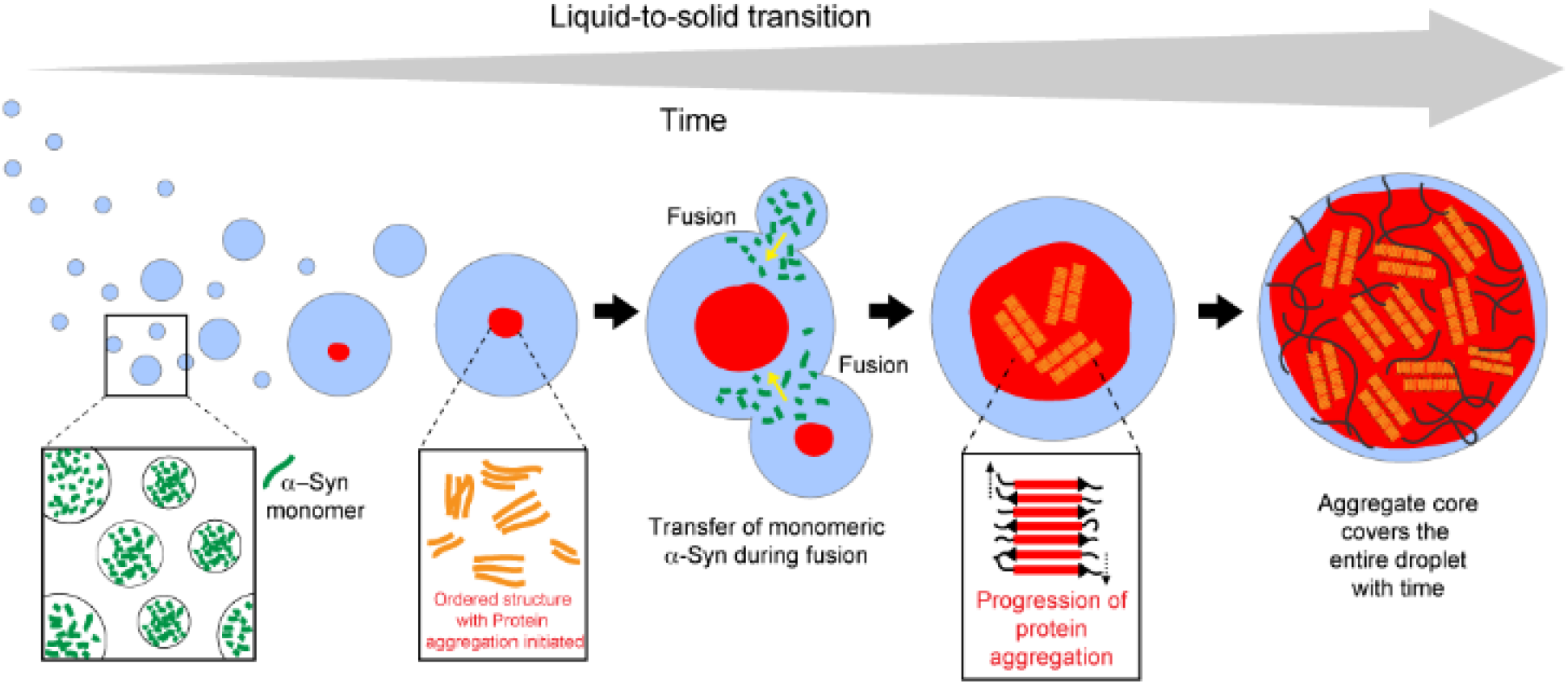
Liquid-liquid phase separation, growth and liquid to solid transition through solid-like core formation at the center of α-Syn droplets. The newly formed phase separated α-Syn droplets contain monomeric protein. During ageing, a solid-like core appears at the center of α-Syn liquid droplets. This core contains ordered structure of α-Syn and gradually converts into β-sheet amyloid with time. The solid-like core expands towards the periphery eventually covering the entire droplet. The growth of the solid-like core (and protein aggregation) is mediated by fusion events. During fusion, smaller, unstable droplets fuse with larger, stable droplets, which already have a structured core. The transfer of monomeric α-Syn from the smaller droplet to the larger droplet (denoted by yellow arrows) further helps in the progression of growth and solidification of the larger droplets.

## Supplementary Information

The Supplementary information file contains materials and methods, Supplementary Fig. 1-11, Supplementary Tables 1-3, Supplementary video legends 1 and 22 references.

## AUTHOR INFORMATION

## Author Contributions

S.R, D.C, K.P, S.M, J.M, A.S.S, M.P, S. K and R.K performed the experiments. All authors participated in analyzing data. S.R and D.C contributed equally^†^ to this work. The study was conceived by S.K.M and designed by S.K.M, G.K, R.P, and A.C. All authors participated in the manuscript writing and approved the manuscript.

## Data availability statement

The authors declare that all the data supporting the findings of this study are available within the paper and in supplementary information files. All the data analysis was performed using published tools and packages and has been cited in the paper and supplementary information text.

## ACKNOWLEDGMENT

We acknowledge IIT Bombay central facilities for TEM, FLIM, FTIRM, TCSPC and Confocal microscopy experiments. We thank Jayant B. Udgaonkar for allowing us to utilize FLIM facility. We thank Sreemantee Sen and Lokesh Ranjan for their help in FLIM experiments.

## ABBREVIATIONS

LLPS: Liquid-liquid phase separation
IDRs: Intrinsically disorder regions
LCDs: Low complexity domains
FLIM: Fluorescence lifetime imaging
FTIRM: Single point Fourier transform infra-red microscopy
FRET: Förster resonance energy transfer
FRAP: Fluorescence recovery after photobleaching
C: Center
IC: Inner Circle
OC: Outer circle
P: Periphery

## Supplementary materials and methods

### Reagents and chemicals

All the reagents and chemicals used in the following experiments were purchased from ThermoFisher Scientific (USA), Sigma (USA), and HiMedia (India) unless otherwise mentioned. Fluorescein-5-maleimide, rhodamine-C2-maleimide and NHS-rhodamine dyes were purchased from ThermoFisher Scientific (USA).

### Expression and purification of recombinant α-Syn

Protein expression and purification was performed using previously established protocol^1–3^. Briefly, *E.coli* BL21-DE3 cells were transformed with pRK172 plasmids encoding for WT and 74C-α-Syn under inducible *lac*-promoter. The expression of the protein(s) was induced by the addition of 1 mM isopropyl-β-D-thiogalactoside (IPTG) to the bacterial culture. After that, the cells were harvested by centrifuging the culture at 4000 rpm for 30 min at 4 °C. The cell pellet is dissolved in appropriate volume of α-Syn buffer—50 mM Tris, pH 8.0, 10 mM EDTA, 150 mM NaCl along with protease inhibitor cocktail (Roche) and subsequently lysed with the help of a probe sonicator (Sonics and Materials Inc. USA) at 4 °C. For 74C-α-Syn, sonication and further steps were performed under a reducing environment in the presence of 1 mM dithiothreitol (DTT) to prevent intermolecular di-sulfide linkages. Next, the cell lysate was subjected to heat denaturation on a boiling water bath for 20 min and centrifuged at 9000 rpm for 30 min at 4 °C. 10% (w/v) streptomycin sulphate and glacial acetic acid was added to the supernatant to precipitate the nucleic acid (DNA) contaminants. The solution is then centrifuged at 9000 rpm for 30 min at 4 °C to pellet the DNA and the supernatant is collected. Equal volume of ice-cold, saturated ammonium sulphate and ultrapure (99%), molecular grade ethanol was added to the supernatant and the solution was kept at 4 °C for ∼12 h to precipitate the protein. The solution is centrifuged at 10000 rpm for 40 min at 4 °C to pellet the protein. The protein pellet was subsequently washed with 100 mM ammonium acetate for 3 to 4 times and precipitated using equal volume of ultrapure ethanol. The protein was finally dissolved in minimum volume of 100 mM ammonium acetate and lyophilized. The lyophilized protein powder was kept at -80 °C for future use. Before and during experiments, the protein quality is checked using SDS-PAGE and MALDI-TOF mass spectrometry (**Supplementary Fig. 1-2**).

### Preparation of aggregate free low molecular weight (LMW) α-Syn

Low molecular weight (LMW) α-Syn was prepared using a standard protocol^3–5^. Briefly, lyophilized protein was dissolved in 20 mM phosphate buffer (pH 7.4), 0.05% sodium azide, and solubilized by adding few drops of 0.2 N (NaOH). The pH of the protein solution was adjusted to 7.4 with 2M HCl with the help of a micro-pH meter probe (Mettler Toledo, USA). The solution was centrifuged at 12000 rpm for 30 min at 4 °C to remove any insoluble impurities. The supernatant was collected and dialyzed against 20 mM phosphate buffer (pH 7.4), 0.05% sodium azide, for ∼12 h using 10 kDa cut off membranes (Sigma, USA). The dialyzed protein solution was then passed through a 100 kDa pre-washed cut-off filter (Merck Millipore, USA) to remove higher order structures (if any). The flow-through contained mostly monomeric α-Syn and is termed as the low molecular weight (LMW) fraction. The concentration of the LMW was determined by measuring the ultraviolet (UV) absorbance at 280 nm. The molar extinction coefficient (ε) of both WT and 74C-α-Syn used for determining concentration was of (5960 M^-1^ cm^-1^)^6^.

### Fluorescence labeling of α-Syn

74C-α-Syn was labeled with fluorescein-5-maleimide at the 74^th^ cysteine residue (at the nonamyloid β component (NAC) region) for fluorescence lifetime imaging (FLIM) experiments. 74C-α-Syn was separately labeled with both fluorescein-5-maleimide and rhodamine-C2-maleimide at the 74^th^ cysteine residue for spectrally (and spatially) resolved Forster resonance energy transfer (FRET) experiments. WT α-Syn was labeled with NHS-rhodamine for fluorescence recovery after photobleaching (FRAP) experiments. All the labeling was performed as per the manufacturer’s (ThermoFisher Scientific, USA) instructions. Briefly, 500 μM of LMW α-Syn (WT and 74C) was mixed with 5 molar excess fluorescent dyes (dissolved in dimethyl sulfoxide (DMSO)). The solution was gently mixed at 25 °C for 2-3 h. Excess or unbound dye was removed by dialysis against 20 mm phosphate buffer (pH 7.4) at 4^0^ C for ∼48 hours, with regular exchange of the buffer at every 5 h interval. The labeled protein concentration and the extent of labeling were measured using a UV spectrophotometer according to the formula provided by the manufacturer. For further experiments, mixture of 10% v/v labeled and 90% v/v unlabeled α-Syn is used to minimize the effect of labeling on phase separation^3^. Our previous study showed that the presence of 10% labeled protein does not affect the structure and aggregation property of WT α-Syn protein^3^.

### *In vitro* liquid-liquid phase separation (LLPS) of α-Syn

LLPS study was performed according to our previously established protocol^3^. Briefly, LMW α-Syn solution containing 200 μM α-Syn and 10% (w/v) PEG-8000 in 20 mM sodium phosphate buffer (pH 7.4), 0.05% sodium azide, was prepared and used for LLPS. Similar protocol was also used for fluorescent-labeled protein as α-Syn shows identical phase behavior irrespective of fluorescent labeling^3^. The LLPS solution (10-15 μl) was then spotted onto a clean glass slide and sandwiched with an 18 mm coverslip (Blue Star, India). The coverslip was then sealed with commercially available nail-polish. The slides were kept in a moist chamber to prevent drying of the LLPS solution and incubated at 37 °C to initiate LLPS. At regular intervals, the slides were taken out and observed under a DMi8 microscope (Leica Microsystems, Germany) at 63X magnification under differential interference contrast (DIC) mode to check for the formation of liquid droplets. In parallel, identical LLPS samples were also incubated in 2 ml eppendorf tubes and at regular intervals, 4-5 μl of sample was drop-casted on a clean glass slide and visualized under 63X magnification under differential interference contrast (DIC) mode to check for the formation of liquid droplets. The details of sample preparation and microscopic observation for fluorescence based studies (FLIM, FRET, FRAP) and Fourier transform infrared microscopy (FTIRM) are given in the respective sections. To check for possible drying of samples, 20 μl of the LLPS samples (200 μM α-Syn + 10% (w/v) PEG-8000) were drop-casted onto 6 mm depression slides and incubated in a moist chamber at 37 °C. The moisture was replenished every 24 h. After d20, the coverslips from the slides are lifted and 5 μl of α-Syn LLPS sample is pipetted and subsequently diluted 10 times. The samples were heated at ∼90 °C for 20 min to denature the higher order structures and fibrils. From the diluted solutions, the concentration (280 nm absorbance) of α-Syn was measured using UV-spectroscopy in the range of 240-340 nm (**Supplementary Fig 4**).

### Design principle for mapping α-Syn liquid droplets under microscope

The α-Syn liquid droplets are spherical in nature. However, microscopy based imaging techniques focus primarily on one-plane along the Z-axis (depth) at a time. This resulted in projection of a spherical droplet as a circular structure during image acquisition. For our microscopy based FLIM and FTIRM experiments; at first, the droplets were imaged at their central plane along the Z-axis, as it is expected to give sufficient information in two dimensions (X-Y plane). The central plane of the droplet was divided into three concentric circles. The radius of the inner most circle (inner circle; IC) was taken as “*r*” and the radii of the next two circles (outer circle: OC and peripheral circle: P) were taken as multiples of “*r*” (2*r* and 3*r*, respectively). The data-points were chosen at the center (C) of the droplet and on the circumference of each concentric circle. From a single droplet, the number of data-points acquired at C, IC, OC and P were 1, 6, 12 and 4, respectively; for fluorescence lifetime imaging (FLIM). On the other hand, the number of data-points acquired at C, IC, OC and P were 1, 6, 6 and 4, respectively; for Fourier transform infra-red microscopy (FTIRM) studies. To qualitatively see whether the pattern observed at the central plane (X-Y axis) was also consistent along the depth (Z-axis); the droplets were subsequently imaged at three equally spaced planes along the Z-axis. The plane closest to the central plane (plane 3) was chosen at a distance *r/3*. The next two planes (plane 2 and plane 1) were taken at distances *2r/3* and *r*, respectively where *r ≈* radius of the droplet. This indicates that the plane 1 chosen was very close to the periphery of the droplet along Z-axis. However, for FTIRM studies, only the central plane could be analyzed (by visually adjusting the focal plane at the largest observable size) because of the instrumental limitations (no depth scanning could be performed).

### Fluorescence lifetime imaging (FLIM) of α-Syn liquid droplets

200 μM of fluorescein-5-maleimide labeled (10% v/v labeled and 90% v/v unlabeled) 74C-α-Syn was phase-separated in presence of 10% w/v PEG-8000 (in 20 mM sodium phosphate buffer, pH 7.4 with 0.05% sodium azide) at 37 °C. The LLPS solution (10 μl) was drop-casted onto a clean glass slide and sandwiched with an 18 mm coverslip (Blue Star, India). The edge of the coverslip was sealed with commercially available nail-polish. These slides were used for subsequent FLIM imaging at different time-points. Furthermore, identical samples were also incubated in 2 ml eppendorf tubes and subjected to FLIM imaging by drop-casting onto clean glass slides just before measurements (**Supplementary Fig. 5**). FLIM was chosen because the technique is able to detect differences in fluorescence lifetime during protein aggregation and can provide a microscopic image of the sample with a spectrum of color coded pixels^7^. The spectrum of the color coded pixels indicates mean lifetime (τ_m_) values (blue being the lowest τ_m_ and red being the highest τ_m_). Moreover, fluorescein was chosen for our experiment as it shows concentration dependent changes of the intensity^8^, which was expected to help delineating the differences in τ_m_ across different regions of phase-separated α-Syn droplets. FLIM images of droplets were obtained at regular time intervals (d2-d20) with a time-resolved microscopy setup (MT-200, PicoQuant; Germany). The phase-separated samples were excited with a 482 nm laser (5-8 µW) pulsing at 40 MHz. From the FLIM images, the fluorescence intensity decay profiles at different regions (C, IC OC and P) of the droplets were recorded from individual pixels under “single-point” recording mode. The fluorescence intensity decay was detected with a dual-band dichroic 480/645 (Chroma) in epifluorescence mode. The droplets were imaged at various Z-planes (planes 1à3) to obtain information along the depth (Z-axis) as mentioned earlier. All the FLIM images and data acquisition was performed using the in-built SymPhoTime 64 software (PicoQuant, Germany).

### Analysis of mean fluorescence lifetime

Fluorescence intensity decay curves were fit with sum of exponential decay equations^7, 9^;

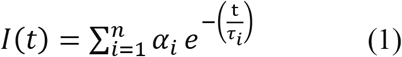

Where *I (t)* = emission fluorescence intensity at magic angle (54.7°) at time t, *α_i_* = amplitude of the fluorescence lifetime given that there are *i* number of lifetimes, *τ _i_* = *i^th^* fluorescence lifetime where, Ʃ*α_i_* = 1.

The fluorescence intensity decay curves obtained from the droplets were satisfactorily fit with biexponential equation^7, 9^ using SymPhoTime 64 software (PicoQuant, Germany). Therefore, for our experiment, equation 1 can be represented as;

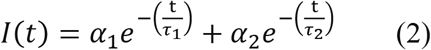

From equation 2, the mean fluorescence lifetime (*τ_m_*) was determined using the following equation^10^

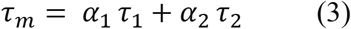

Where,

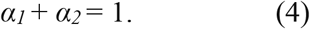

For each sample, the mean long (Λ_m_) fluorescence lifetime and mean short (λ_m_) fluorescence lifetime were computed. Since we did not know the origin of the long and short lifetimes, the overall mean fluorescence lifetime (τ_m_) was calculated from each of the data-points on the droplets as a relative measure of our observations. Each region was denoted with different color codes (▪) with respect to its τ_m_. The color codes were then assigned to the respective data-points to generate a spatially resolved map (SRM) of the liquid droplets.

For two fusing droplets at d5, multiple data-points were chosen on the FLIM image across the longitudinal fusion axis (p1→p6). Although fusion is a dynamic process, the timescale of fusion (minutes) was much larger than the timescale of FLIM image acquisition (seconds)—enabling it to be experimentally feasible. From each of these data-points, the respective fluorescence intensity decay was obtained. The decays were subsequently fit to bi-exponential function to compute the τ_m_ values. All the experimental and instrument parameters were identical and the data analyses were performed as described in the previous section.

### FTIRM imaging of α-Syn liquid droplets

Fourier transform infra-red microscopy (FTIRM) was performed with unlabeled WT α-Syn protein. 200 μM of unlabeled α-Syn was mixed with 10% (w/v) PEG-8000 (in 20 mM sodium phosphate buffer, pH 7.4, with 0.05% sodium azide) and incubated at 37 °C to initiate LLPS. The LLPS solution was drop-casted onto a clean glass slide (Blue Star, India) and the formation of liquid droplets was confirmed by visualizing the LLPS solution under a DMi8 microscope (Leica Microsystems, Germany) in DIC mode. At different time intervals (d2-d20) during liquid-to-solid transition, the solution was spotted on a BaF_2_ coated glass slide (38 x 19 x 4 mm) (Technosearch Instruments, India). BaF_2_ coated slide was used since it shows minimal absorption property in the entire UV-IR spectrum (200 nm to 12 µm) and is resistant towards high energy radiation. The experiment was performed in the presence of a Vertex-80v vacuum optics bench (Bruker, Germany), to minimize the presence of atmospheric water during FTIR spectrum acquisition. The liquid droplets were visualized and imaged at their central plane by manually focusing the droplet image until the largest diameter (size) is achieved by a 3000 Hyperion microscope attached with Vertex-80 FTIR machine (Bruker, Germany) at 20X magnification (ATR objective lens—0.5 µm resolution). Individual data-points were assigned on the FTIRM image of the droplets at different locations (C-Center, IC-Inner Circle, OC-Outer Circle and P-Periphery). However, due to the instrumental limitations, images along the Z-plane could not be obtained for FTIRM technique. Due to the smaller size of the liquid droplet (2-3 μm) at d2, the data-points were obtained from all the pixels representing a single liquid droplet. FTIR spectra were recorded in “single-point mapping” mode^11, 12^ in the wavenumber range of 1500-1800 cm^-1^. This corresponds to the amide-I (C=O stretching) and amide-II (C-N stretching & N-H bending) regions of a peptide bond showing predominant absorption intensity in that wavenumber range. Notably, the widely used “focal plane array” (FPA) mode^11–13^ could not be used for our experiments because the droplet size was very close to the resolution limit of the FTIRM (0.5 µm) at earlier time-points. Therefore, to maximize the signal to noise ratio, single-point mapping mode was chosen for our study.

### FTIR spectra analysis

FTIR spectra analysis was done with OPUS-65 v6.5 software. For each data point, background correction was implemented to eliminate the contributions from the solvent (and water) near 1650 cm^-1^. Subsequently, the background corrected spectra were subjected to baseline correction and “Fourier Self Deconvolution” (FSD) in the wavenumber range of 1600-1700 cm^-1^ corresponding to the amide-I (C=O stretching) frequency. This was done because all of the protein secondary structures correspond to amide-I stretching in FTIR^14^. The deconvolution technique was performed to obtain single sharp peaks for each secondary structure which were masked by the convoluted (broadened) spectra. The FSD was performed using the Fourier transformed Lorentzian deconvolution function^15^. Furthermore, the deconvolution was performed by feeding band and noise deconvolution factors^15^ to the OPUS-65 v6.5 software. These factors were optimized to obtain maximum sharpness and minimum noise. After that, every peak maxima corresponding to different secondary structures were assigned on the deconvoluted spectra. The spectra were best fitted by auto-fitting method with minimum residual root mean square (RMS) error. The integrated area under each individual spectrum represented the fractional contribution from corresponding secondary structure. The integration limit was taken from 1600 cm^-1^ to 1700 cm^-1^, since all of the secondary structures appeared within the amide-I stretching frequency^14, 15^. Each of the integration values was divided from the sum of all the integration values and multiplied with 100 to obtain the % abundance of each secondary structure. To schematically represent our findings, we assigned various color codes (represented with ▪; one for each data-point) to denote the prevalent secondary structural combinations at different regions of the droplet with time. To conclusively state our observations, we took a fairly large sample-size (n=15 droplets) for our FTIRM experiment.

To understand the secondary structure of α-Syn at different locations of two fusing droplets (d5), the fusion event was imaged and three data-points (p1, p2 and p3) were assigned along the longitudinal fusion axis. The FTIR spectra were obtained from these data-points and subsequently analyzed as mentioned earlier. This could be possible because the time of acquisition of FTIRM spectra from a fusing droplet was in the time-range of seconds; which was much less than the timescale of fusion event (in minutes).

### *In vitro* fluorescence recovery after photobleaching (FRAP) of α-Syn liquid droplets

200 µM NHS-rhodamine labeled (10% v/v labeled + 90% v/v unlabeled) α-Syn was phase-separated in the presence 10 % PEG-8000 (in 20 mM sodium phosphate buffer, pH 7.4, 0.05% sodium azide). The LLPS solution was drop-casted onto clean glass slides (Blue Star, India) and sandwiched with 18 mm coverslip (Blue Star, India). At two different time-points (d5 and d20) during LLPS, photo-bleaching of the phase separated α-Syn droplets undergoing fusion event was performed using a standard protocol^3^ with a confocal microscope with an in-built FRAP setup (Zeiss Axio-Observer Z1 microscope). Briefly, a 561 nm DPSS 561-10 laser (at 100% laser power) was used to bleach the larger droplet between the two droplets undergoing fusion. The bleaching region (ROI-1, termed as R1) was chosen on the larger droplet. Another ROI (termed as R2) of the same radius (ω) was chosen on the smaller droplet. The post-bleach fluorescence intensity was recorded from both R1 and R2, simultaneously. The passive bleaching by the monitoring laser was recorded from a different droplet (not undergoing fusion). The background correction was performed from the droplet free region. All the experimental parameters were identical and data analyses were performed as described in the following section. All of the microscopic snapshots were obtained with a 1024 x 1024 frame size with 8 bit-depth.

For spatially resolved FRAP experiments at different location in the same droplet; d5, d7 and d15 LLPS samples (incubated in 2 ml eppendorf tubes) were spotted onto clean glass slides (Blue Star, India) and immediately imaged under a confocal microscope with 63x oil immersion objective (Zeiss Axio-Observer Z1 microscope). The bleaching regions R1, R2 and R3 were chosen at the C, OC and P regions, respectively and the fluorescence recovery was recorded for a span of 200 s. The laser intensity and relevant instrumental parameters were kept identical for all time-points and for all bleaching regions. The microscopic snapshots were obtained with a 1024 x 1024 frame size with 8 bit-depth.

### FRAP data analysis

Recovery of fluorescence obtained from different ROIs (R1, R2, and R3) were corrected at each time-frame for background and passive bleaching. This was done by computing the normalized fluorescence intensity, *I (n)* using the following equation^3^;

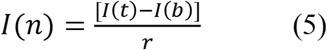

Where, *I (t)* = fluorescence intensity at time t,

*I (b)* = background fluorescence intensity,

*r* = *I_c_*/*I_c_0__* which is the rate of photo bleaching,

Where, *I_c_0__* = fluorescence intensity of the given ROI before photo-bleaching,

*I_c_* = fluorescence intensity of the given ROI after photo-bleaching.

From equation 9, the mobile fraction (M.F or *A*) of the protein molecule in the given ROI can be calculated using the following equation^3^;

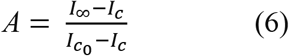

Where, *I*_∞_ = fluorescence intensity at the end of recovery time-period The normalized and background corrected fluorescence recovery was fitted using a single exponential equation with the help of OriginPro 9.0 (OriginLab, USA) software. The equation is as follows^3, 16–20^;

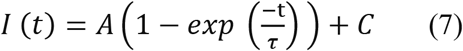

Where, τ = fluorescence recovery time constant,

*A* = mobile fraction of the fluorescent probe,

*C* = y-intercept of the recovery curve (if any).

### Spatially (and spectrally) resolved FRET microscopy

Spectrally resolved FRET microscopy was performed by following a previously established protocol^3, 21^. Briefly, LLPS solution (200 μM protein + 10% PEG-8000) of fluorescein-5-maleimide (Donor, *D*) labeled (10% v/v labeled) 74C-α-Syn was mixed equal volume of LLPS solution of rhodamine-C2-maleimide (Acceptor, *A*) labeled (10% v/v labeled) 74C-α-Syn. The solution was drop-casted on a clean glass-slide and sandwiched with an 18 mm coverslip and sealed for phase separation. Only *D*-labeled 74C-α-Syn and only *A*-labeled 74C-α-Syn was also phase separated simultaneously, which served as *D* only and *A* only controls, respectively.

Spectrally resolved images of the droplets were obtained from both FRET and control samples at d2 (immediately after formation of liquid droplets), d5 (initial stage during liquid-to-solid transition) and d20 (final stage during liquid-to-solid transition). The images were acquired manually at the central plane using an epifluorescence total internal reflection fluorescence microscope (Nikon Eclipse TE2000-U). *D* and *A* were excited with 488 nm and 532 nm DPSS lasers (OXXIUS, model: ACX-CTRB and LASERGLOW, model: LRS-0532-PFM-00200-03), respectively. Single droplets were visualized with a 60X TIRF objective lens (1.49 NA) (Nikon, Japan). The co-localization of *D* and *A* labeled droplet was obtained by emissions at two energetically separated detection channels (515 – 565 nm and 590 − 700 nm band-pass emission channels). The droplets were imaged at 50 ms exposure. A background flattening of each image was performed using ImageJ (Fiji, NIH). Spatially (as well as spectrally) resolved image of the droplets were acquired using a slit and a transmission grating (70 g/mm) using a sCMOS camera (Hamamatsu ORCA-Flash 4.0 V3). The dispersed emission spectra (in the range 500-700 nm) from individual droplets were recorded in this manner. All the images were collected at identical excitation power (5W/cm^2^) of 488 nm and for same emission (at 532 nm) exposure time (0.3 s). All measurements were performed at 25 °C and the data were analyzed using ImageJ (Fiji, NIH) and OriginPro 9.0 (OriginLab, USA).

### FRET microscopy data analyses

The data analysis of single droplet FRET measurement was performed according our previously established protocol^3^. Briefly, the *D* only emission spectra were obtained by exciting the sample at 488 nm (donor (*D*) excitation wavelength). The *A* only droplets showed ∼5.4 times greater fluorescence signal when excited at 532 nm (acceptor (*A*) excitation wavelength) compared to 488 nm excitation. This was due to the difference of the absorption coefficients of *A* at 488 and 532 nm.

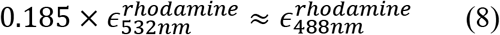

We also noticed a long tail in the emission spectrum of *D*, which was extended beyond the emission maxima of *A* (at 595 nm). Therefore, corrections for the emission tail and the difference in *ϵ* were performed according to our previously established protocol^3^. Subsequently, to calculate the apparent FRET efficiency, we defined a semi-quantitative term called the intensity enhancement factor due to the energy transfer (*IEF_ET_*), which was directly proportional to the actual FRET efficiency. The *IEF_ET_* accounted for both the corrections mentioned above.

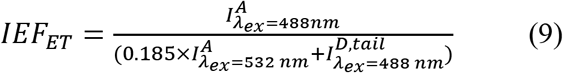

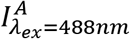 = fluorescence intensity of *A* when excited at 488 nm,

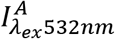 = fluorescence intensity of *A* when excited at 532 nm,

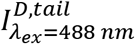 = fluorescence intensity of the *D* emission tail when excited at 488 nm for the FRET samples.

In an actual FRET scenario, we can expect a positive enhancement of *IEF_ET_* (*i.e. IEF_ET_* >1). The instrumental parameters were identical for the FRET samples as well as for *D* only and *A* only controls. The concentrations of the fluorescent dyes were also equal in both controls and FRET samples.

### Electron Microscopy (EM) of α-Syn liquid droplets

TEM imaging was done following a previously established protocol^3^. Briefly, 200 μM of unlabeled α-Syn was mixed with 10% (w/v) PEG-8000 (in 20 mM sodium phosphate buffer, pH 7.4, 0.05% sodium azide) and incubated at 37 °C to initiate LLPS. The LLPS solution was drop-casted onto a clean glass slide (Blue Star, India) and sandwiched with an 18 mm coverslip (Blue Star, India). The formation of liquid droplets was confirmed by visualizing the LLPS solution under a DMi8 microscope (Leica Microsystems, Germany) in DIC mode. At different time-points (d2-d15), the coverslips were removed from the respective slides and the samples were transferred directly on the EM grid (Electron Microscopy Sciences, USA). The grids were negatively stained with aqueous 1% (w/v) uranyl formate solution and subsequently imaged with a transmission electron microscope (Phillips CM-200, Amsterdam, Netherlands). The imaging was performed at 200 KV and 6600X magnification and collected by Keen View Soft imaging system (Olympus, Tokyo, Japan). For scanning electron microscopy (SEM) experiments, the LLPS sample (d5) was dropcasted onto a clean glass coverslip and immediately imaged in cryo-mode with the help of a JSM-7600F SEM (JEOL, Japan).

### TEM image analysis of α-Syn liquid droplets

The obtained TEM images at different time-points were analyzed with the help of ImageJ software (Fiji, NIH). A reference line along the entire diameter of the droplet was drawn on the TEM image and the grayscale intensity along the line was computed. Important to note, the pixel intensity values were higher at brighter regions of the image and lower at darker regions. Subsequently, angular maps^22^ of the droplets at individual time-points were generated by straightening the droplet image from 0 to 180° with the help of an in-built “straighten” module in ImageJ (Fiji, NIH). To conclusively state our observations, a large number of sample-size was chosen for our analyses (n=20 droplets).

## Supplementary Figures

**Supplementary Fig. 1:**
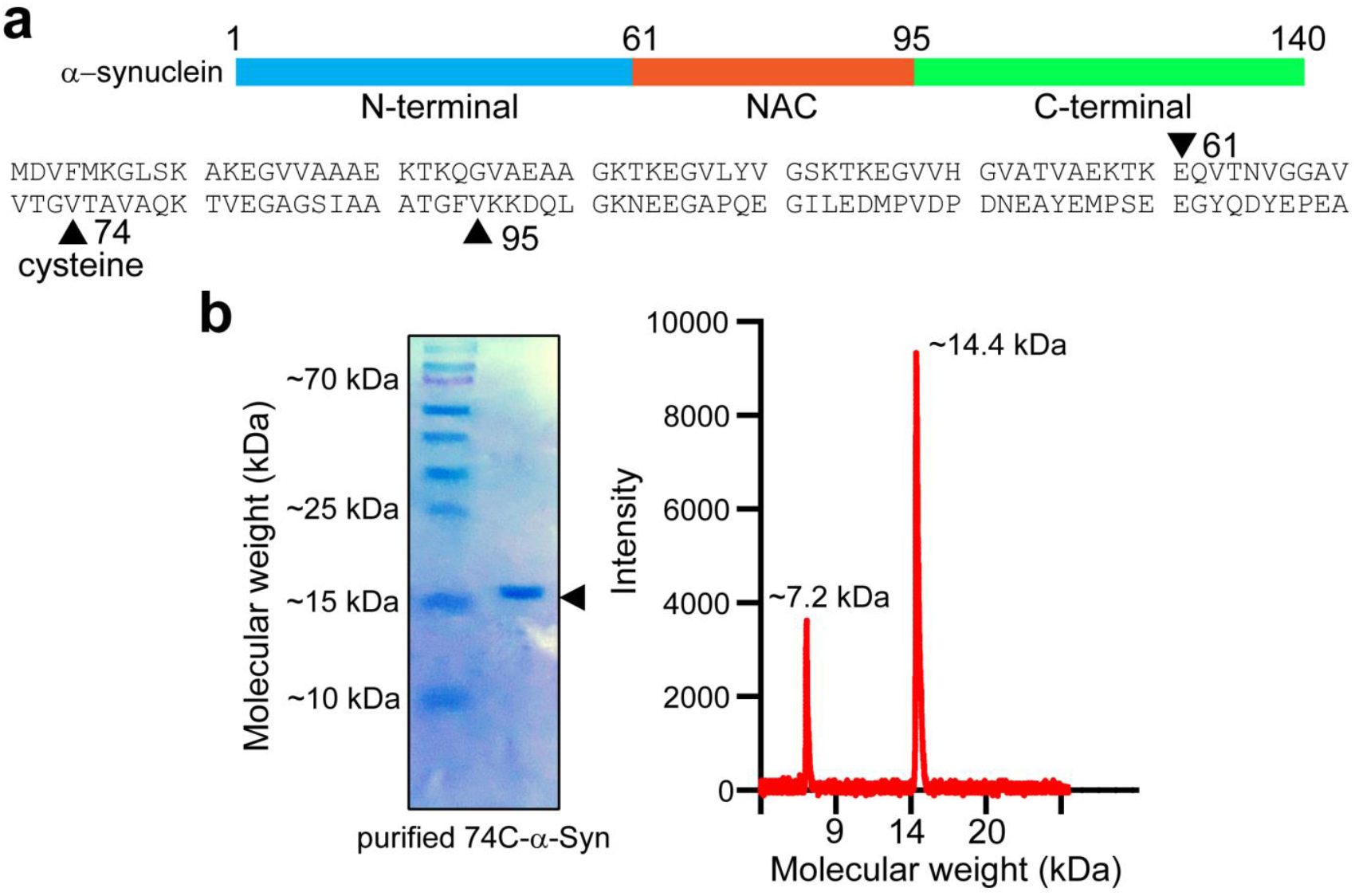
Purification of 74C-α-Syn for FLIM studies. **a. *Upper panel:*** Schematic diagram showing the amphipathic N-terminal (1-60), hydrophobic NAC (61-95) and acidic C-terminal (96-140) domain of wild type (WT) human α-Syn. ***Lower panel:*** The amino acid sequence of full length WT human α-Syn is shown. The start and end of the hydrophobic NAC domain is denoted with black triangular pointers (▲) at residues 61-95, respectively. The 74^th^ valine residue (▲) is mutated with cysteine to get the 74C-α-Syn mutant which has been used for FLIM and FRET studies. **b. *Left panel:*** SDS-PAGE of the purified 74C-α-Syn showing a single band at ∼15 kDa representing the full length protein. ***Right panel:*** Mass spectrometry analysis also confirms that the purified 74C-α-Syn is intact by showing two major peaks (m/z-∼14.4 kDa and m/2z∼7.2 kDa).

**Supplementary Fig. 2:**
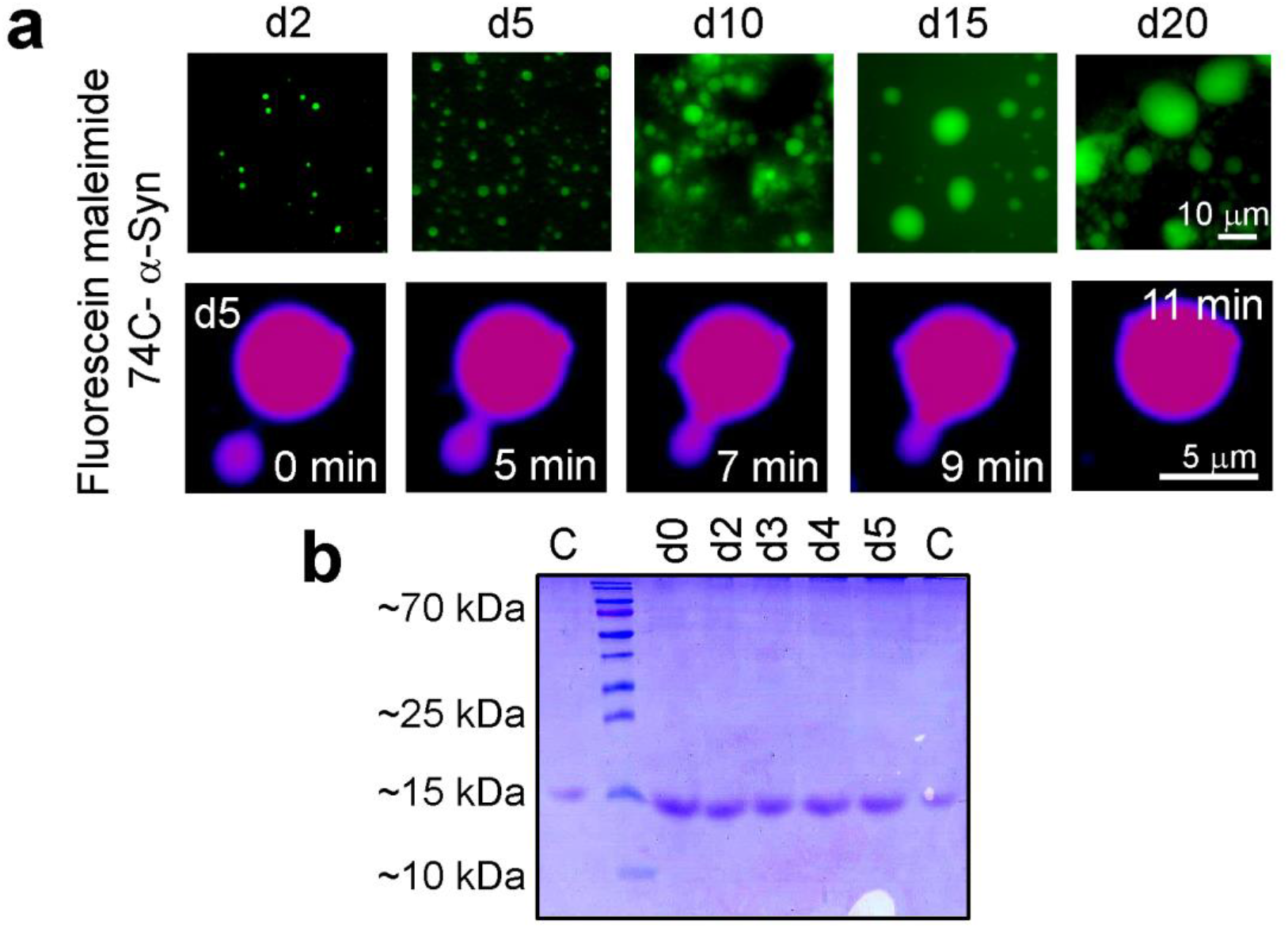
LLPS behavior of labeled α-Syn a. *Upper panel:* Fluorescein-5-maleimide labeled 74C-α-Syn undergoes LLPS in presence of 10% PEG-8000. The droplets continuously grow both in size and number indicating the progression of LLPS. Representative images are shown. n=3 independent observations. ***Lower panel:*** Fusion event (at d5) between two fluorescein maleimide labeled 74C-α-Syn droplets is shown. Representative event is shown. The experiment is carried out two times with similar observations. The images are represented in thermal LUT for better visualization. **b.** SDS-PAGE of 74C-α-Syn at different time points (d0-d5) showing that the protein remains intact during LLPS. “C” represents monomeric 74C-α-Syn, which has been used as a control.

**Supplementary Fig. 3:**
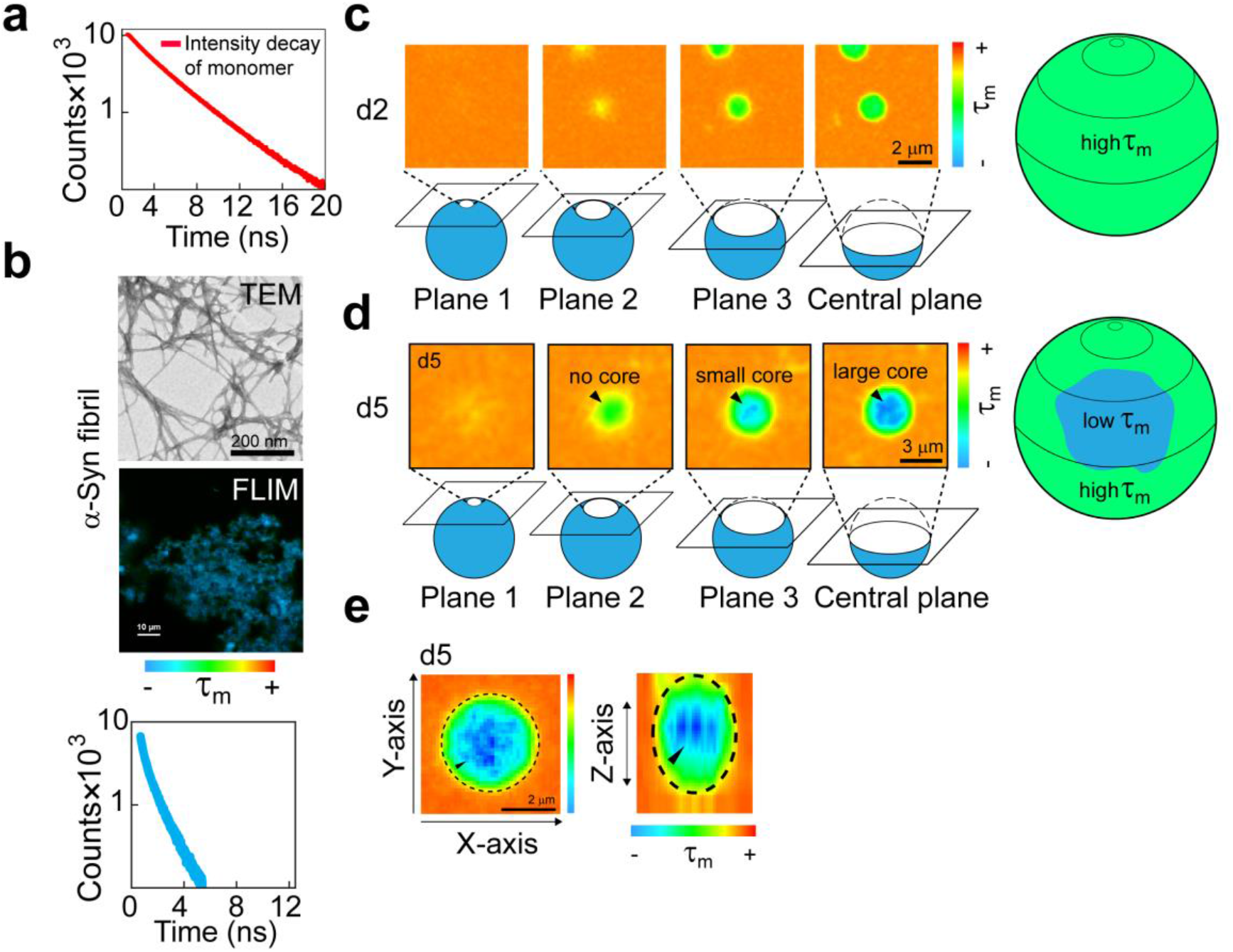
Distribution of the mean fluorescence lifetime (τ_m_) of α-Syn phase-separated droplets during liquid-to-solid transition. **a.** Time-resolved fluorescence intensity decay of soluble, fluorescein-5-maleimide labeled 74C-α-Syn monomer control. The overall mean lifetime (τ_m_) is calculated to be ∼4.1±1 ns. Representative intensity decay profile is shown. The experiment is performed three times with similar results. **b. *Upper panel:*** Transmission electron microscopy (TEM) imaging of independently prepared fluorescein-5-maleimide labeled 74C-α-Syn amyloid fibrils. Representative images are shown. The experiment is performed three times with similar observations. ***Middle panel:*** Corresponding fluorescence lifetime images (FLIM) of the fibrils from which the time-resolved intensity decay is obtained. The color spectrum represents the range of mean lifetime (τ_m_) (blue being the lowest and red being the highest). ***Lower panel:*** Time-resolved fluorescence intensity decay of fluorescein-5-maleimide labeled 74C-α-Syn fibril control. Representative decay profile is shown. The experiment is performed three times with similar results. **c. *Left panel:*** FLIM images of the droplets obtained at different Z-planes from the periphery to the center (plane1→ 3, central plane) showing similar color image (similar τ_m_) of α-Syn at each plane in the d2 droplet. The FLIM image (top) and corresponding plane (bottom, schematically drawn) are shown. The FLIM images were manually brightened to highlight the differences of mean lifetimes across different regions. ***Right panel:*** Schematic diagram of a spherical d2 droplet showing similar life time (τ_m_) inside the droplet at different planes. **d. *Left panel:*** Fluorescence lifetime (FLIM) image of d5 droplets at different Z-planes (plane 1, 2, 3 and central plane) shows reduced τ_m_ (blue color, denoted with black triangular pointers) when the Z-plane is close to the central plane (plane 3). For planes 1 and 2, which are away from the central plane (i.e. away from the center of the droplet), τ_m_ becomes higher (green color). ***Right panel:*** Schematic diagram of a spherical d5 droplet showing the formation of a core structure at the center of the droplet, which has a different local microenvironment (reduced τ_m_). The images are manually brightened to highlight the differences of mean lifetimes across different regions. **e.** FLIM images of planes 1→3 (along with the central plane) are superimposed. Orthogonal (Z) projection of the superimposed image is shown along the Z-axis to show the spherical nature of the droplet. This data also confirms that the reduced τ_m_ is indeed at the center (indicated by black triangular pointer (▴)) of a spherical, d5 liquid droplet. The images are manually brightened to highlight the differences of mean lifetimes across different regions. The black dashed line represents the boundary of the droplet. The experiment is performed three times with similar observations.

**Supplementary Figure 4:**
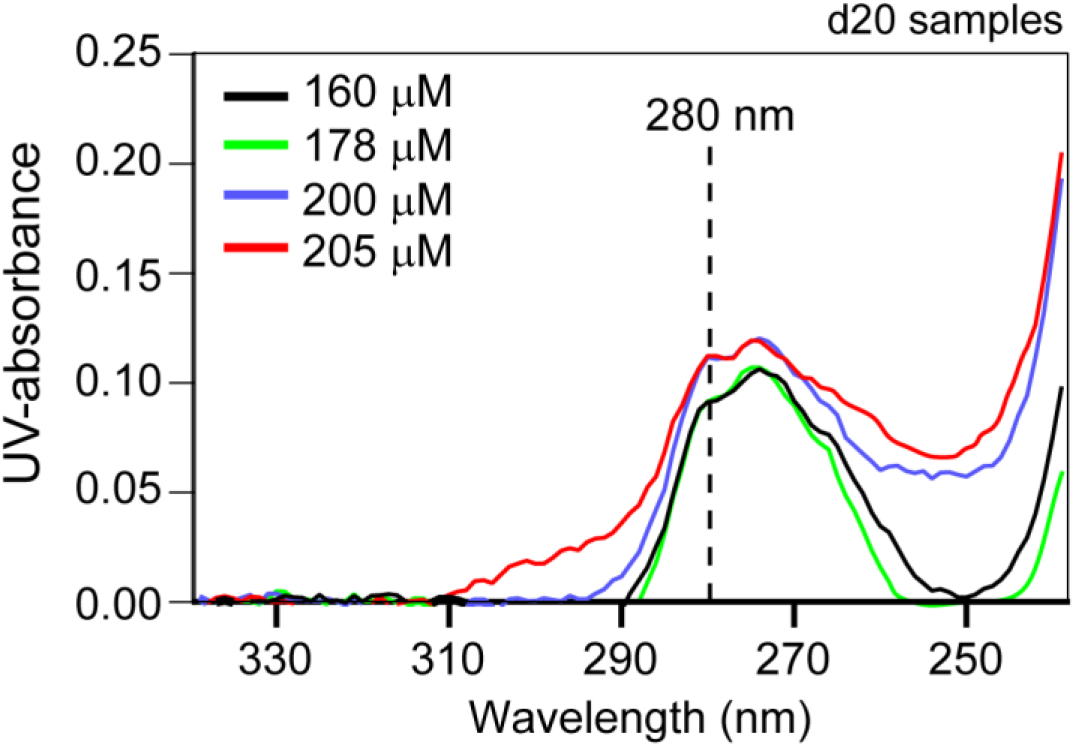
Incubation of LLPS sample on a slide does not lead to drying when kept in a moist chamber with frequent replenishment of moisture. UV-Visible spectra of four different samples diluted from drop casted liquid droplet sample showing >80% proteins are retained in the sample even after 20 days of incubation. This suggests that even after d20, the concentration of the LLPS samples do not change drastically from the original 200 μM concentration. Even though the LLPS sample is heated at 90 °C for several minutes to denature any higher order aggregates, the hydrogel-like nature of the system after d20 may lead to inhomogeneous partitioning of the α-Syn molecules while pipetting. Therefore, slight deviation (such as in the case of 160 µM and 178 µM) from the original concentration may be observed.

**Supplementary Figure 5:**
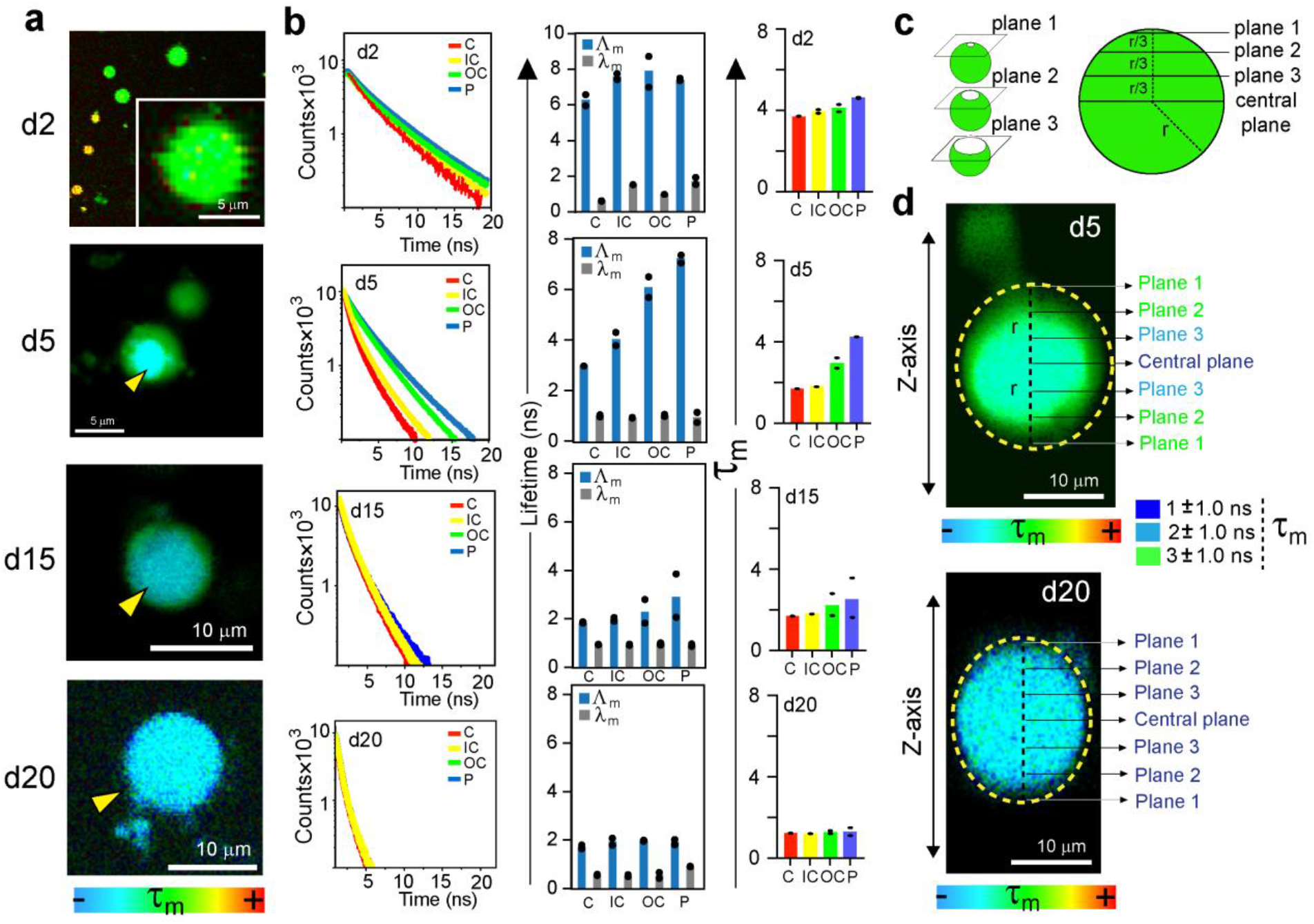
Changes in fluorescence lifetime initiates at the center and progresses towards the periphery of α-Syn phase-separated droplets during liquid-to-solid transition when the LLPS samples are incubated in tubes. **a**. Representative color coded original FLIM images showing fluorescein-5-maleimide labeled 74C-α-Syn droplets at various time points (d2-d20) during liquid-to-solid transition. The color spectrum represents the range of overall mean lifetime (τ_m_) (blue being the lowest and red being the highest). The images are acquired from the central plane. The yellow triangles mark the solid-like core at d5, d15 and d20. **b.** (Left panel) Time resolved fluorescence intensity decay obtained from different regions (C-Center, IC-Inner Circle, OC-Outer circle and P-Periphery) of the droplets at various time points (d2-d20). Representative spectra are shown. The experiment is performed two times with similar observations. (Middle panel) The long (Δ_m_) and short (λ_m_) lifetimes obtained from the intensity decay curved are plotted for C-Center, IC-Inner circle, OC-Outer circle and P-Periphery at different time-points (d2, d5, d15 and d20). (Right panel) Corresponding mean lifetimes (τ_m_) obtained from intensity decay curve at different regions (C-Center, IC-Inner circle, OC-Outer circle and P-Periphery) of the phase-separated droplets are plotted for different time-points (d2, d5, d15 and d20). **c.** Schematic showing analysis of droplets along the depth (Z-axis). Apart from the central plane, three other planes (plane 1, 2 and 3) along the Z-axis of the droplets are also imaged for FLIM to examine the distribution of lifetimes along the depth. The planes are equally spaced with a distance of *r/3* with radius of the droplet is *r*. **d. (**Upper panel) FLIM image of a d5 droplet along the Z-axis is shown. The mean lifetimes (τ_m_) calculated from different planes along the Z-axis confirms the presence of solid-like core at the center of the phase separated droplet. (Lower panel) FLIM image of a d20 droplet along the Z-axis is shown. The mean lifetimes (τ_m_) calculated from different planes along the Z-axis indicates the homogeneous nature of the droplet with a reduced τ_m_ indicating the progression of solid-like core to the periphery of the droplet. The experiment is performed two independent times with similar observations.

**Supplementary Figure 6:**
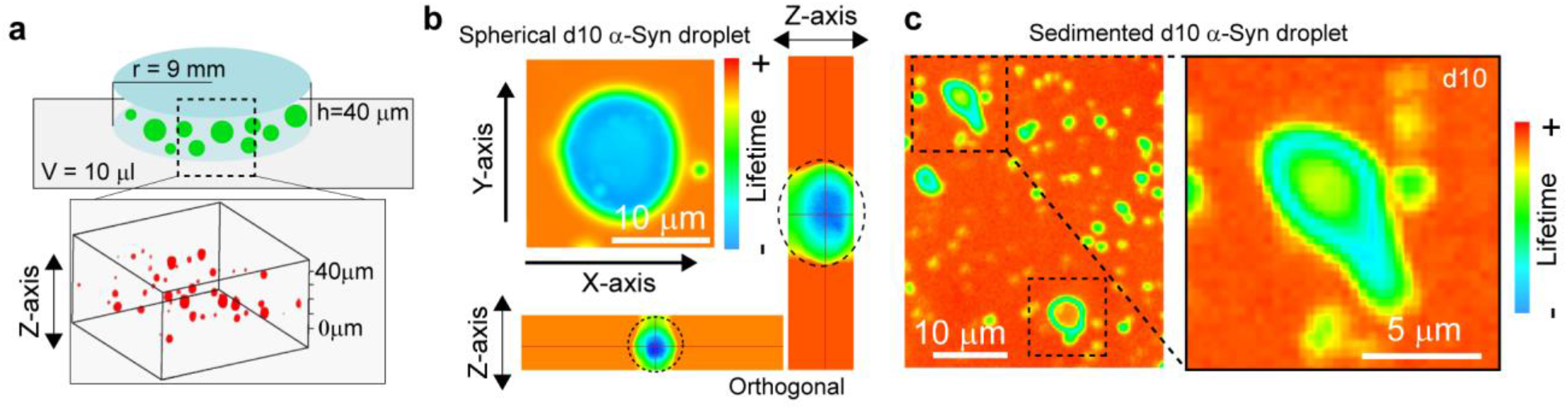
Analysis of α-Syn phase separated droplets that are geometrically spherical. **a.** Schematic representation of the drop-casted phase separated sample, which creates a cylinder when sandwiched with a coverslip with a radius (r) of 9 mm. The depth (h) of the cylinder is calculated to be ∼40 μm. **Bottom panel** represents a 3d-reconstitution of a real Z-scan data obtained from the microscope, which shows spherical d5 α-Syn droplets in the solution. **b.** FLIM depth scanning (along Z-axis) of a d10 droplet containing a solid-like core (reduced lifetime-indicated with blue color) clearly shows the spherical nature of the droplet. **c.** At d10, some of the droplets sediment at the bottom of the glass slide and lose their spherical shape. **Inset** represents a d10 droplet showing a deformed lifetime distribution as well as geometry when it is in contact with the glass surface. Representative images are shown.

**Supplementary Fig. 7:**
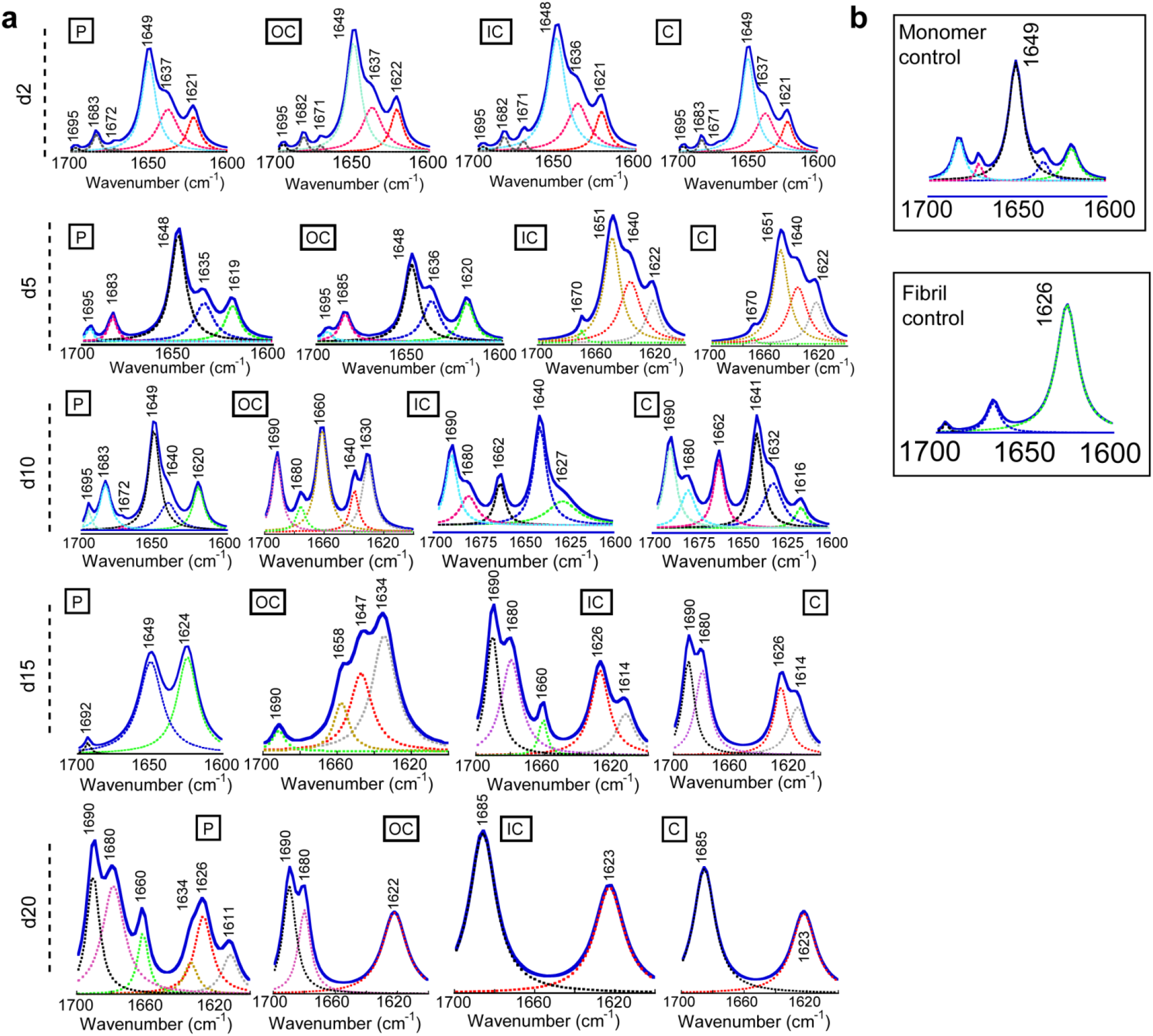
Fourier-transform infrared (FTIR) spectra of various regions of α-Syn phase-separated droplet during liquid-to-solid transition. **a.** Representative FTIR spectra obtained from different regions (C-Center, IC-Inner Circle, OC-Outer circle and P-Periphery) of the microscopic image of α-Syn droplets at d2, d5, d10, d15 and d20 are shown. **b. (*Boxed*)**: FTIR spectra of α-Syn monomer (***upper panel***) and pre-formed α-Syn fibril (***lower panel***) are shown as controls. The individual peaks corresponding to different secondary structures are assigned (dotted lines) on the deconvoluted spectra (solid blue line) **(a-b)**. The experiment is performed three times with similar results.

**Supplementary Fig. 8:**
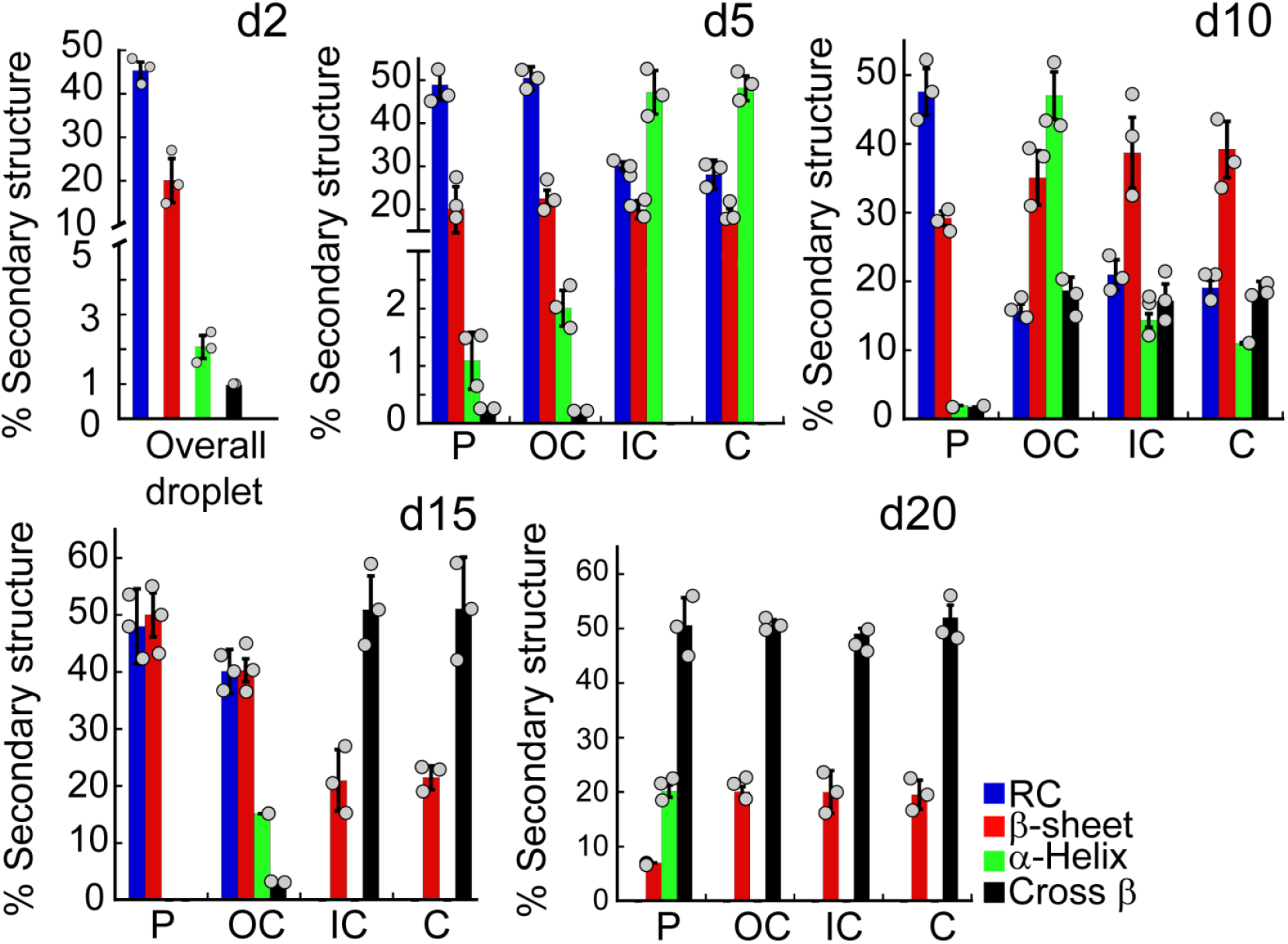
Conformational transition of α-Syn inside the phase-separated droplets. FTIR spectra are deconvoluted and the % of the secondary structure for different regions (C-Center, IC-Inner circle, OC-Outer circle and P-Periphery) on the droplets at various time-points (d2, d5, d10, d15 and d20) are determined. The % of random coil (RC), β-sheet, α-helix and cross-β are plotted. Values represent mean ± S.D for n=3 independent experiments.

**Supplementary Fig. 9:**
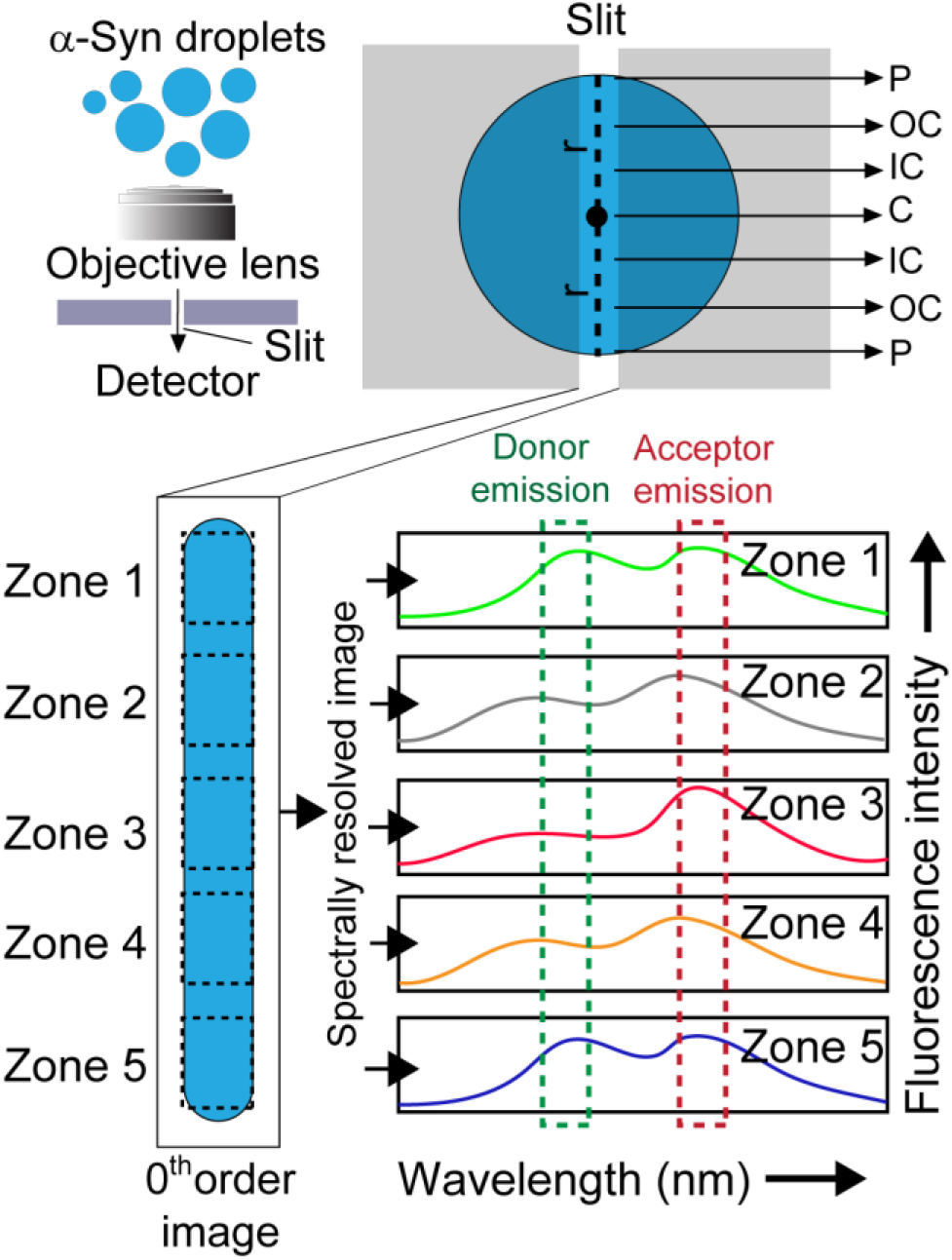
Working principle behind spatially resolved FRET microscopy of α-Syn LLPS. Schematic diagram of the working principle for FRET microscopy is shown. ***Top left:*** Phase separated α-Syn droplets labeled with both fluorescein-5-maleimide (Donor, *D*) and rhodamine-C2-maleimide (Acceptor, *A*) labeled 74C-α-Syn, which is visualized under a coverslip through a narrow slit. ***Top right:*** The slit is positioned in such a way so that the droplet is visualized along its entire diameter (Radius (***r***) ***+ r***). Different regions (C-Center, IC-Inner Circle, OC-Outer circle and P-Periphery) have been assigned along the diameter of the droplet. ***Bottom left***: The microscopic image obtained by this alignment is called the 0^th^ order image^3, 21^. The 0^th^ order image along the diameter can be imaged with the excitation wavelength of *D* and emission wavelengths of both *D* and *A*, in a location specific (Zones 1-5) manner. By doing this, we can generate a spatially as well as spectrally resolved image^3^ of the droplet. The fluorescence intensity profiles at different emission wavelengths for different locations (Zones 1-5) **(*Bottom right)*** enable us to calculate the intensity enhancement factor (*IEF_ET_*) which is a proportional measure of the FRET efficiency^3^.

**Supplementary Fig. 10:**
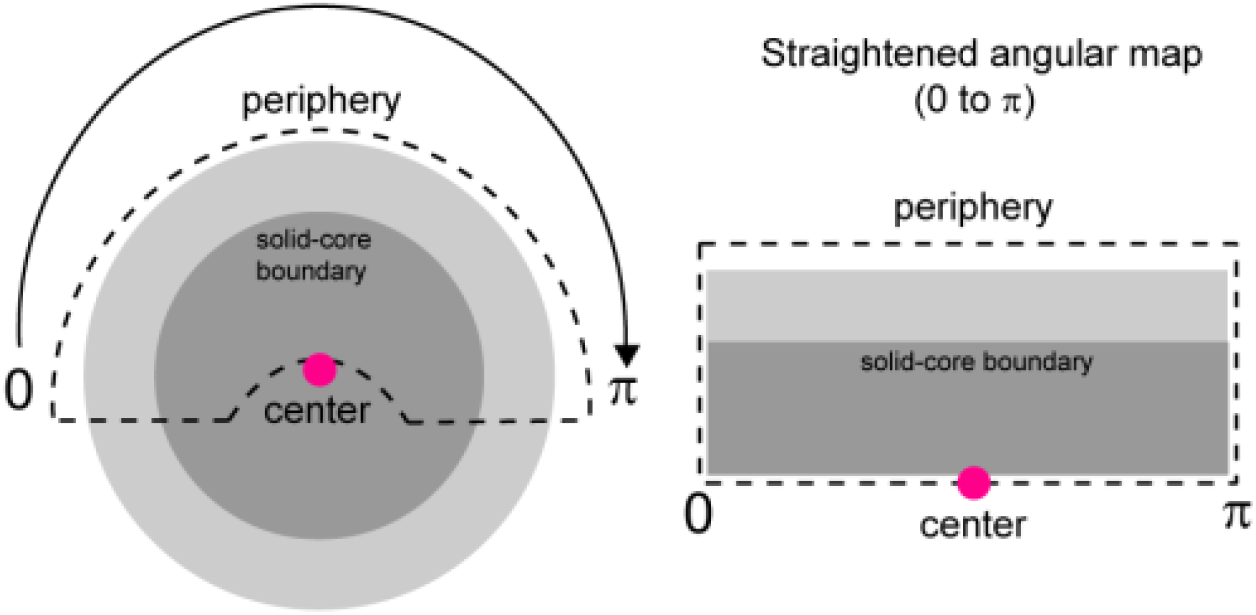
Angular maps of TEM images of α-Syn liquid droplets. ***Left panel:*** Schematic representing a phase-separated α-Syn droplet (light gray) containing a solid-like core (dark gray) observed under TEM. The black dashed boundary denotes the area (spanning 0◊π or 0°◊180° angle of the droplet) from which the angular map (***right panel***) has been generated^22^. The center of the droplet is marked with a pink dot (●). ***Right panel:*** The straightened, angular map (0◊π or 0°◊180° angle) of the droplet has been schematically represented. The center of the droplet is marked with a pink dot (●). From here, the distance of the solid-like core boundary from the center is easily quantified with respect to the radius.

**Supplementary Figure 11:**
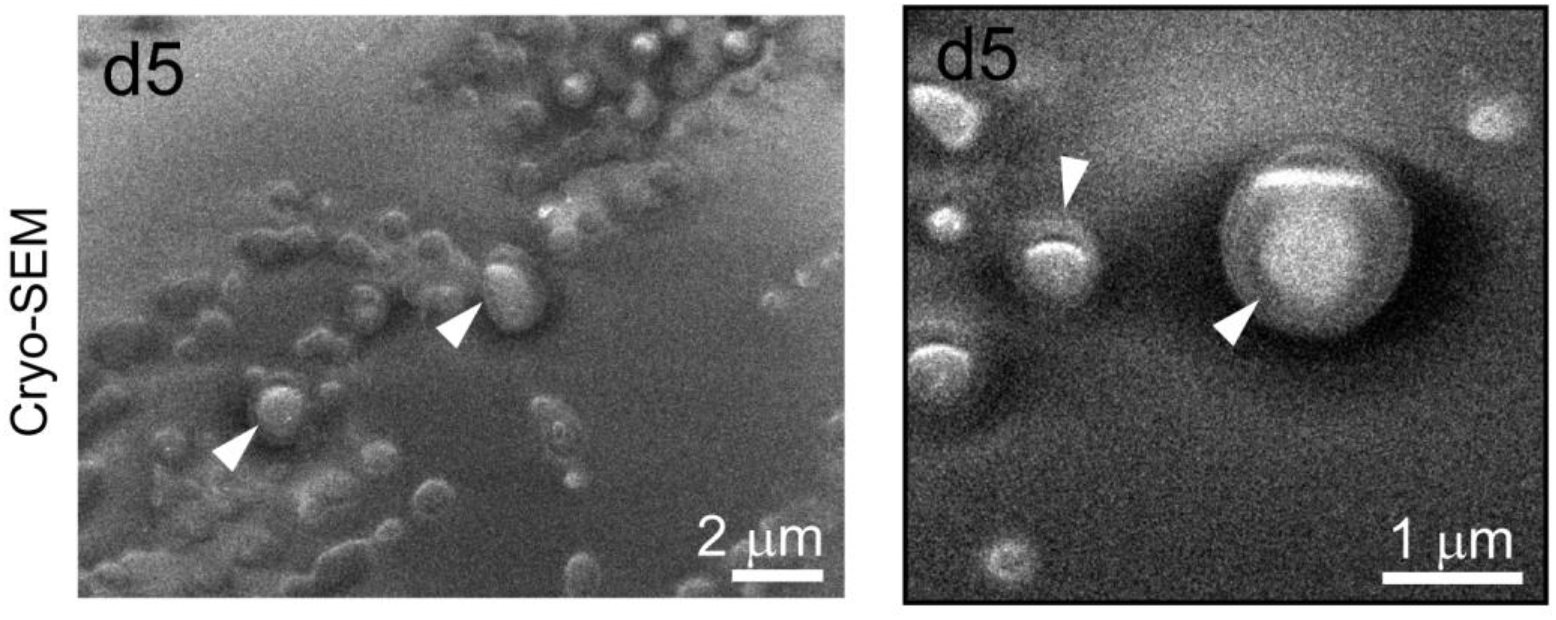
Cryo-SEM imaging of α-Syn liquid droplets at d5. Cryo-SEM imaging of d5 droplets showing the presence of a core structure (marked with white triangular pointers) supporting TEM observations of solid like dense structure at the center of liquid droplet. Representative images from two different experiments are shown.

**Supplementary Table 1:** Deconvoluted amide-I stretching frequencies in the range of 1600-1700 cm^-1^ and their corresponding protein secondary structures^14^.

| Secondary Structure | Start wavenumber ($\text{cm}^{-1}$) | End wavenumber ($\text{cm}^{-1}$) |
| --- | --- | --- |
| Random coil (RC) | $1640 \pm 2$ | $1648 \pm 2$ |
| β-sheet | $1620 \pm 2$ | $1638 \pm 2$ |
| α-helix | $1650 \pm 2$ | $1660 \pm 2$ |
| Cross-β amyloid | $1690 \pm 2$ | $1695 \pm 2$ |

**Supplementary Table 2:** The % of the secondary structures of α-Syn at different locations (C-Center, IC-Inner circle, OC-Outer circle and P-Periphery) of the phase-separated droplets during liquid-to-solid transition (d5-d20). The % of turn and loop structures are not mentioned in the table.

| Time (days) | Secondary structural conformation | Percentage of different secondary structures at different location inside the droplet |  |  |  |
| --- | --- | --- | --- | --- | --- |
|  |  | P (%) | OC (%) | IC (%) | C (%) |
| d5 | RC | 48.9 | 50.4 | 30.0 | 28.0 |
| | β-sheet | 20.1 | 22.4 | 20.0 | 19.1 |
| | α-helix | 1.1 | 2.0 | 47.2 | 48.1 |
| | Cross-β | 0.3 | 0.2 | ~0 | ~0 |
| d10 | RC | 47.5 | 15.6 | 20.9 | 19.1 |
| | β-sheet | 29.2 | 35.1 | 38.7 | 39.2 |
| | α-helix | 1.9 | 47.0 | 14.3 | 11.0 |
| | Cross-β | 1.8 | 18.7 | 17.1 | 18.0 |
| d15 | RC | 48.0 | 40.1 | ~0 | ~0 |
| | β-sheet | 50.0 | 40.3 | 20.9 | 21.4 |
| | α-helix | ~0 | 15.1 | ~0 | ~0 |
| | Cross-β | ~0 | 3.1 | 50.9 | 51.0 |
| d20 | RC | ~0 | ~0 | ~0 | ~0 |
| | β-sheet | 7.0 | 20.0 | 20.0 | 19.5 |
| | α-helix | 20.1 | ~0 | ~0 | ~0 |
| | Cross-β | 50.5 | 50.5 | 49.0 | 52.0 |

**Supplementary Table 3:**
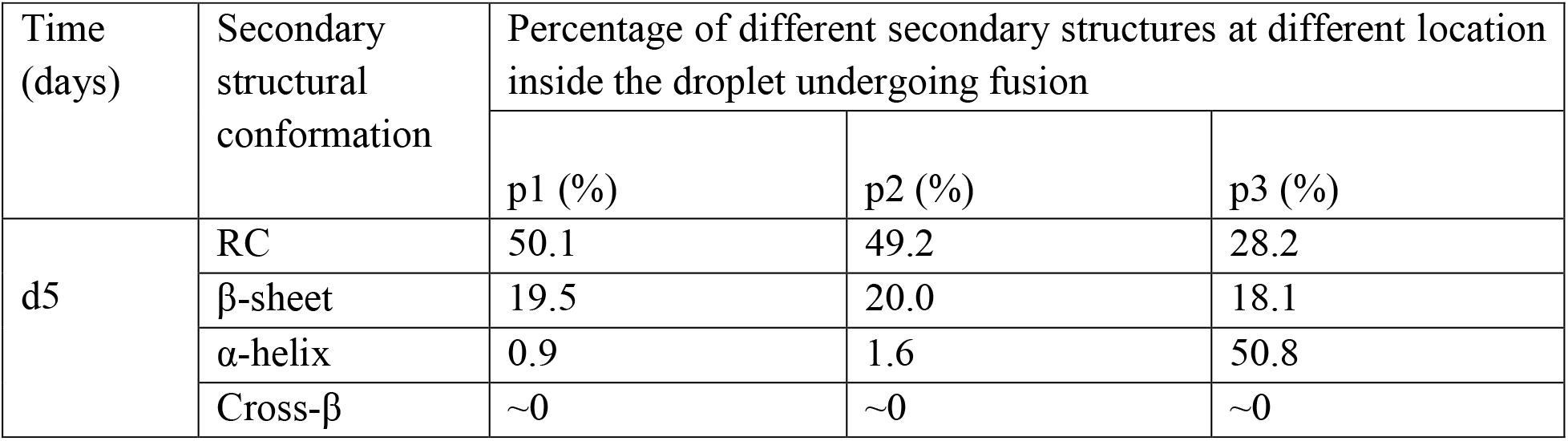
The % of the secondary structures of α-Syn at different locations (p1-contact region of fusion, p2-center of smaller droplet and p3-center of larger droplet) of the phase-separated droplets undergoing fusion event at d5. The % of turn and loop structures are not mentioned in the table.

| Time (days) | Secondary structural conformation | Percentage of different secondary structures at different location inside the droplet undergoing fusion |  |  |
| --- | --- | --- | --- | --- |
|  |  | p1 (%) | p2 (%) | p3 (%) |
| d5 | RC | 50.1 | 49.2 | 28.2 |
| | β-sheet | 19.5 | 20.0 | 18.1 |
| | α-helix | 0.9 | 1.6 | 50.8 |
| | Cross-β | ~0 | ~0 | ~0 |

## Supplementary video legends

**Supplementary video 1:** Spatially resolved FRAP of d5 α-Syn droplet confirm that the molecules at the center are less diffusive compared to the periphery due to the formation of a solid-like core.

